# Transcript profiling of *plastid ferrochelatase two* mutants reveals that chloroplast singlet oxygen signals lead to global changes in RNA profiles and are mediated by Plant U-Box 4

**DOI:** 10.1101/2024.05.13.593788

**Authors:** Snigdha Rai, Matthew D. Lemke, Anika M. Arias, Maria F. Gomez Mendez, Katayoon Dehesh, Jesse D. Woodson

**Affiliations:** The School of Plant Sciences, University of Arizona, Tucson, AZ; Department of Botany and Plant Sciences, Institute for Integrative Genome Biology, University of California, Riverside, CA

**Keywords:** Abiotic stress, *Arabidopsis thaliana*, chloroplast, programmed cell death, reactive oxygen species, singlet oxygen, jasmonic acid, salicylic acid

## Abstract

**Background:** In response to environmental stresses, chloroplasts generate reactive oxygen species, including singlet oxygen (^1^O_2_), an excited state of oxygen that regulates chloroplast-to-nucleus (retrograde) signaling, chloroplast turnover, and programmed cell death (PCD). Yet, the central signaling mechanisms and downstream responses remain poorly understood. The *Arabidopsis thaliana plastid ferrochelatase two* (*fc2*) mutant conditionally accumulates ^1^O_2_ and Plant U-Box 4 (PUB4), a cytoplasmic E3 ubiquitin ligase, is involved in propagating ^1^O_2_ signals for chloroplast turnover and cellular degradation. Thus, the *fc2* and *fc2 pub4* mutants are useful genetic tools to elucidate these signaling pathways. Previous studies have focused on the role of ^1^O_2_ in promoting cellular degradation in *fc2* mutants, but its impact on retrograde signaling from mature chloroplasts (the major site of ^1^O_2_ production) is poorly understood.

**Results:** To gain mechanistic insights into ^1^O_2_ signaling pathways, we compared transcriptomes of adult wt, *fc2*, and *fc2 pub4* plants. The accumulation of ^1^O_2_ in *fc2* plants broadly repressed genes involved in chloroplast function and photosynthesis, while inducing genes and transcription factors involved in abiotic and biotic stress, the biosynthesis of jasmonic acid (JA) and salicylic acid (SA), microautophagy, and senescence. Elevated JA and SA levels were observed in ^1^O_2_-stressed *fc2* plants. *pub4* reversed most of this ^1^O_2_-induced gene expression and reduced the JA content in *fc2* plants. The *pub4* mutation also blocked JA-induced senescence pathways in the dark. However*, fc2 pub4 plants* maintained constitutively elevated levels of SA even in the absence of bulk ^1^O_2_ accumulation.

**Conclusions:** Together, this work demonstrates that in *fc2* plants, ^1^O_2_ leads to a robust retrograde signal that may protect cells by downregulating photosynthesis and ROS production while simultaneously mounting a stress response involving SA and JA. The induction of microautophagy and senescence pathways indicate that ^1^O_2_-induced cellular degradation is a genetic response to this stress, and the bulk of this transcriptional response is modulated by the PUB4 protein. However, the effect of *pub4* on hormone synthesis and signaling is complex and indicates that an intricate interplay of SA and JA are involved in promoting stress responses and programmed cell death during photo-oxidative damage.

## Background

Plants, as sessile organisms, are constantly exposed to a variety of environmental challenges that can severely influence their growth, development, and ultimately, survival. These challenges (i.e., abiotic, and biotic stresses) trigger multilevel responses that involve stress sensing, signal transduction, regulation of gene expression, and post-translational protein modifications. As sites of photosynthesis, chloroplasts are sensitive to environmental fluctuations, and can act as stress sensors. These organelles produce reactive oxygen species (ROS), including superoxide and hydrogen peroxide at photosystem I (PSI) and singlet oxygen (^1^O_2_) at photosystem II [1]. This ROS production is exacerbated under stresses such as high or excess light (EL) [2], drought [3], heat [4], and salinity [5]. ROS is highly reactive and can damage cellular macromolecules, including lipids, proteins, and DNA. If unchecked, ROS can impair chloroplast function, disrupt photosynthesis, and lead to cell death. In photosynthetic tissues, ^1^O_2_ is the predominant ROS, causing most of the photo-oxidative damage [2]. However, ^1^O_2_ can also serve as a stress signaling metabolite, leading to the regulation of nuclear gene expression (via plastid-to-nucleus retrograde signaling), chloroplast degradation, and programmed cell death (PCD) [6–8]. As ^1^O_2_ has a short half-life (∼1 µsec) and limited diffusion distance (∼200 nm) [9], signaling pathways likely employ secondary messengers that exit the chloroplast to elicit a signal. However, the molecular mechanisms by which these signals act are largely unknown.

One challenge to studying ^1^O_2_ signaling in plant cells, is that chloroplast ^1^O_2_ is produced simultaneously with other types of ROS in response to natural stresses like EL [7]. Thus, it can be difficult to attribute specific responses to ^1^O_2_ during photo-oxidative stress. Researchers have turned to *Arabidopsis thaliana* mutants that conditionally accumulate chloroplast ^1^O_2_ to avoid this complication. One mutant, *plastid ferrochelatase 2* (*fc2*), is defective in one of two conserved plastid ferrochelatases (FC2) that produces heme from protoporphyrin IX (ProtoIX) in the chloroplast-localized tetrapyrrole (i.e., chlorophylls and hemes) biosynthesis pathway [10].

The tetrapyrrole pathway is activated at dawn, leading to an increase in ProtoIX synthesis within chloroplasts [11]. In *fc2* mutants grown under diurnal cycling light conditions, ProtoIX accumulates, becomes photo-sensitized, and quickly produces ^1^O_2_ [12, 13]. This results in differential regulation of nuclear gene expression, chloroplast degradation, and eventually PCD. Under permissive constant light conditions, *fc2* mutants avoid PCD, but still accumulate a small amount of ProtoIX and ^1^O_2_ [14]. This leads to the selective degradation of a subset of chloroplasts. Up to 29% of chloroplasts are in some stage of degradation in a single *fc2* cell, and 8% (or about 1/3 of degrading chloroplasts) interact with the central vacuole and generate “blebbing” structures that push into the vacuole compartment [14]. This process was demonstrated to be independent of autophagosome formation [15] and may instead be facilitated by fission-type microautophagy, a poorly understood process in eukaryotic cells [16]. Envelope proteins in ^1^O_2_-damaged chloroplasts are also ubiquitinated prior to chloroplast degradation and the initiation of PCD [12, 17], which may be a mechanism cells use to target damaged chloroplasts for turnover. Such a chloroplast quality control pathway would ensure healthy populations of chloroplasts performing efficient photosynthesis [18]. As 80% of leaf nitrogen is stored within these organelles, they are also prime targets for nutrient redistribution [19]. The induction of PCD may also be a means to avoid necrosis and water loss [7].

Forward genetic screens in *fc2* [12, 20] and other ^1^O_2_-producing mutants [21] have isolated mutations that block these signals, demonstrating that ^1^O_2_-induced cellular degradation is genetically controlled. The cytoplasmic E3 ubiquitin ligase Plant U Box 4 (PUB4) was identified in this way as playing a signaling role [12]. *fc2-1 pub4-6* mutants (referred to as *fc2 pub4* hereafter) showed elevated levels of ProtoIX and ^1^O_2_ compared to wt plants, yet the chloroplasts were not ubiquitinated or degraded, nor was PCD initiated [12, 20]. This has led to a model where PUB4 is involved in a ^1^O_2_ signal to control chloroplast turnover and PCD, possibly by ubiquitinating proteins on damaged chloroplasts and targeting them for degradation and/or by modulating chloroplast-to-nucleus retrograde signaling [6]. Further studies have shown that PUB4 is involved in chloroplast degradation and PCD induced by EL and other sources of photo-oxidative stress [17, 20, 22, 23] and PUB4 has been shown to associate with degrading chloroplasts [24]. However, the mechanism by which this occurs is not known and no PUB4 in vivo ubiquitination targets have been identified.

In addition to cellular degradation pathways, chloroplast ^1^O_2_ can lead to chloroplast-to-nucleus retrograde signaling and affect the expression of hundreds of genes [25]. To understand the impact of ^1^O_2_ in *fc2*, we previously performed a microarray analysis of de-etiolating (greening) *fc2* seedlings [12], which demonstrated that these mutants experience photo-oxidative damage, induce putative fission-type microautophagy genes [15], and their RNA profiles resemble other ^1^O_2_ over accumulating mutants [22]. However, because this analysis used etiolated seedlings, it is not clear what impact ^1^O_2_ will have on mature chloroplasts in photosynthetic tissue and true leaves, which are the main sites of ^1^O_2_ production in plants [2]. Furthermore, no genetic suppressors of *fc2* were included in this analysis to indicate which ^1^O_2_ signaling pathways were activated.

To further understand the molecular mechanisms underlying ^1^O_2_ stress signaling and the role of PUB4 in photosynthetic tissue, here we have measured global RNA profiles of *fc2* and *fc2 pub4* plants in ^1^O_2_-producing cycling light conditions. We demonstrate that ^1^O_2_ from mature chloroplasts leads to significant alterations to global transcript profiles in adult *fc2* plants, particularly affecting chloroplast and photosynthesis-associated genes. *pub4* largely reversed these effects, restoring transcript levels to those observed in wt plants. ^1^O_2_ also led to the induction of jasmonic acid (JA) and salicylic acid (SA) biosynthesis, senescence pathways, and microautophagy. *pub4* reversed these effects except for the accumulation of SA, which was constitutive in *fc2 pub4* plants, indicating a link between PUB4 and SA metabolism and signaling. *pub4* was also shown to block dark- and JA-induced senescence, indicating that PUB4 may play a general role in senescence pathways triggered by photo-oxidative stress. Together, our results demonstrate that chloroplast ^1^O_2_ signaling is mediated by PUB4 and broadly regulates nuclear gene expression and downstream responses, thereby playing an important role in regulating a cell’s response to photo-oxidative stress.

## Results

### The singlet oxygen stress response in adult stage *fc2* mutants is blocked by the *pub4* mutation

Under constant (24h) light conditions, Arabidopsis *fc2* plants appear healthy and do not experience PCD. However, when grown under diurnal light cycling conditions, they accumulate ^1^O_2_, which triggers retrograde signaling, bulk chloroplast degradation, and eventually PCD [12, 14, 15]. This phenomenon can be observed in adult plants where necrotic lesions appear on leaves [20, 22]. On the other hand, *fc2 pub4* mutants exhibit elevated levels of ^1^O_2_ compared to wt, yet they do not undergo PCD or chloroplast degradation [12, 20]. The molecular mechanism behind ^1^O_2_-induced signaling in *fc2* and its reversal by *pub4* is unknown. To understand this, we scrutinized these mutants in the adult stage under permissive 24h light and ^1^O_2_-generating 16h light/8h dark diurnal (cycling) conditions. This allowed us to assess the impact of ^1^O_2_ in mature chloroplasts, which is the major site of ^1^O_2_ production in plants [2].

In 24h light conditions, four-week-old wt, *fc2*, and *fc2 pub4* plants appear healthy (**Figs. 1a and b**). However, when grown for three weeks in 24h light conditions before transitioning to cycling light conditions for one week, *fc2* exhibits leaf lesions, whereas *fc2 pub4* does not. Trypan blue stains dead cells and was used to quantify cell death (**Figs. 1c-e**). As expected, *fc2* showed a significant increase in cell death compared to wt under two and seven days of cycling light conditions. This was blocked by the *pub4* mutation and *fc2 pub4* showed a level of cell viability similar to wt.

**Figure 1:**
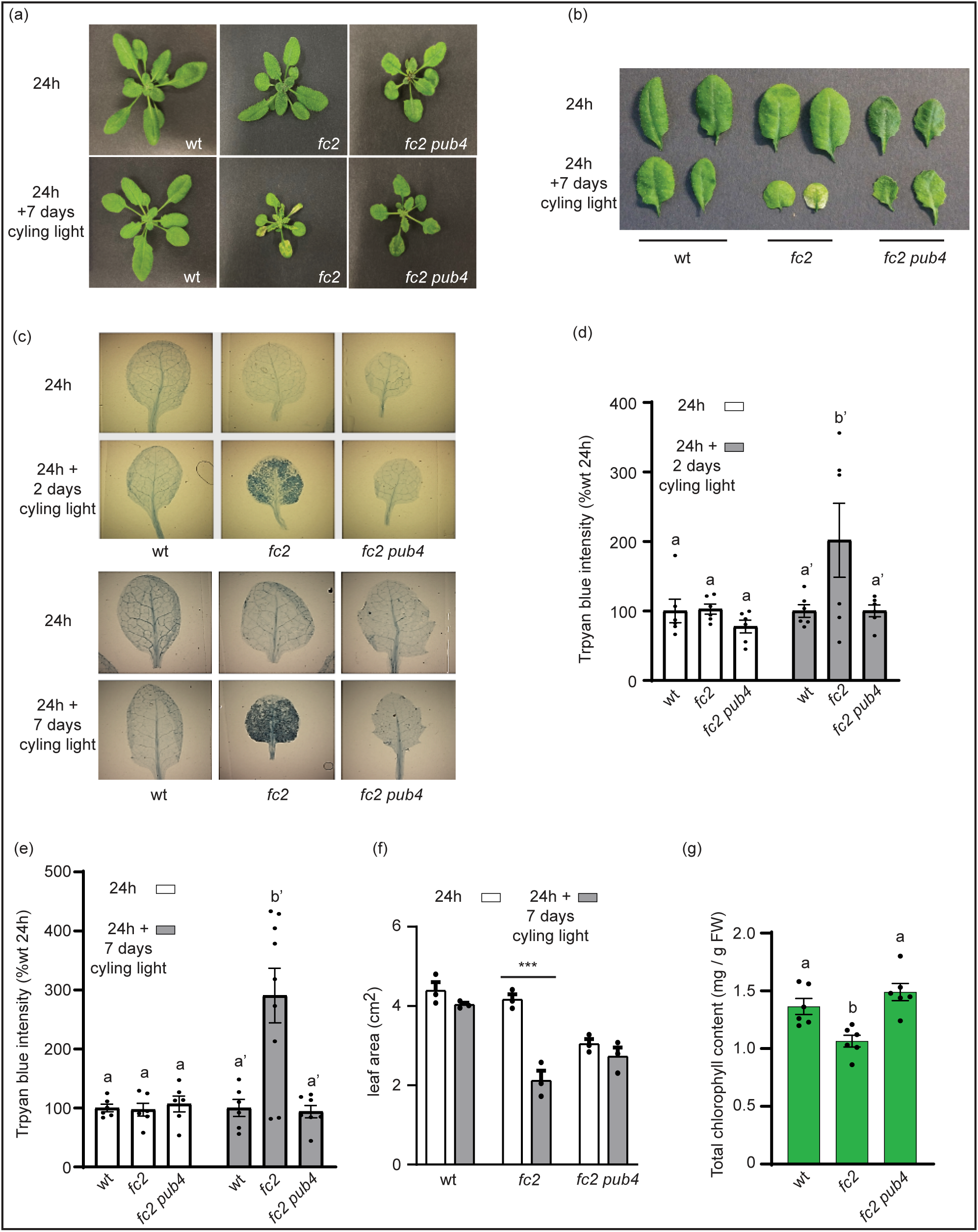
The singlet oxygen stress response in *fc2* adult plants is blocked by the *pub4* mutation. The effect of chloroplast ^1^O_2_ accumulation was assessed in adult plants. (a) Four-week-old plants grown in constant (24h) light or under stressed conditions (three weeks in 24h light and one week in 16h light/8h dark diurnal (cycling) light conditions). (b) Representative rosette leaves (#’s 3 and 4) from the plants in (a). (c) Representative leaves (#’s 3 and 4) of plants after two and seven days of cycling light conditions, stained with trypan blue. The deep dark color is indicative of cell death. (d) and (e) show mean values (± SEM) of the trypan blue signal in leaves shown in (c) (n ≥ 6 leaves from separate plants). (f) shows the mean total leaf area (±SEM) of plants in panel (a) (n = 3 plants). (g) Mean total chlorophyll content (mg/g FW) (+/− SEM) of adult plant leaves grown in 24h light conditions (n = 6 biological replicates). For panels d, e, and g, statistical analyses were performed by a one-way ANOVA test followed by Tukey’s HSD. Letters indicate statistically significant differences between samples (*P* ≤ 0.05). For panel f, a two-way ANOVA followed by Sidak multiple comparison test were used for statistical analysis (n = 3 leaves from separate plants) (***, *P* value ≤ 0.001). Closed circles represent individual data points.

Under 24h light conditions, *fc2* mutants exhibit pale rosettes in comparison to wt, albeit without a significant change in leaf area (**Figs. 1a, b, and f**). On the other hand, *fc2 pub4* mutants are green, but have significantly reduced leaf area (̴ 30%) compared to wt. Consistently, *fc2* has reduced total chlorophyll content compared to wt, and this effect is reversed by *pub4* (**Fig. 1g**). When transferred to cycling light conditions, *fc2* exhibits impaired growth, with 50% reduced leaf area compared to wt (**Figs. 1a, b, and f**). Conversely, *fc2 pub4* plants do not exhibit any significant growth reduction after a change in light conditions. Taken together, our results show that the *pub4* mutation mitigates the negative impact of chloroplast ^1^O_2_ stress in adult stage *fc2* mutants.

### The *pub4* mutation alters the transcriptional responses to singlet oxygen stress

To gain insight into how ^1^O_2_ may impact the *fc2* mutant, and what effect the *pub4* mutation has on this response, we performed a whole transcriptome analysis of wt, *fc2*, and *fc2 pub4* plants. Plants were first grown for 16 days in permissive 24h light conditions. To initiate ^1^O_2_ stress, they were then shifted to cycling light conditions for two days and leaves were harvested one hour after subjective dawn (cycling light sets). A second set of plants remained in 24h light conditions for the full 18 days (24h sets). A Principal Component Analysis (PCA) analysis (**Fig. S1a**) and distance matrix plot (**Fig. S1b**) were performed with the replicate samples (three each for a total of 18), indicating that the light condition had the largest effect on global gene expression and that there was less clustering of genotypes under cycling light conditions compared to 24h light conditions. A pairwise comparison of gene expression between the mutants and wt supported this conclusion (**Figs. S2a and b**), with more differences observed in the cycling light conditions irrespective of genotype.

To identify genes controlled by ^1^O_2_ and/or PUB4, we performed four pairwise comparisons between wt and the mutants in permissive 24h light (“*fc2* vs. wt - 24h”, *fc2* compared to wt, grown in 24h light; “*fc2 pub4* vs. wt - 24h”, *fc2 pub4* compared to wt grown under 24h constant light conditions) or ^1^O_2_-inducing cycling light conditions (“*fc2* vs. wt - cycling”, *fc2* compared to wt grown in cycling light conditions; “*fc2 pub4* vs. wt - cycling”, *fc2 pub4* compared to wt grown under cycling light conditions). We then applied cutoffs (Log_2_ Fold-Change (FC) > +/− 1.5, *padj* < 0.01) to identify a total of 1682 differentially expressed genes (DEGs) within the dataset (**Fig. 2A** and **Tables S1-5**).

**Figure 2:**
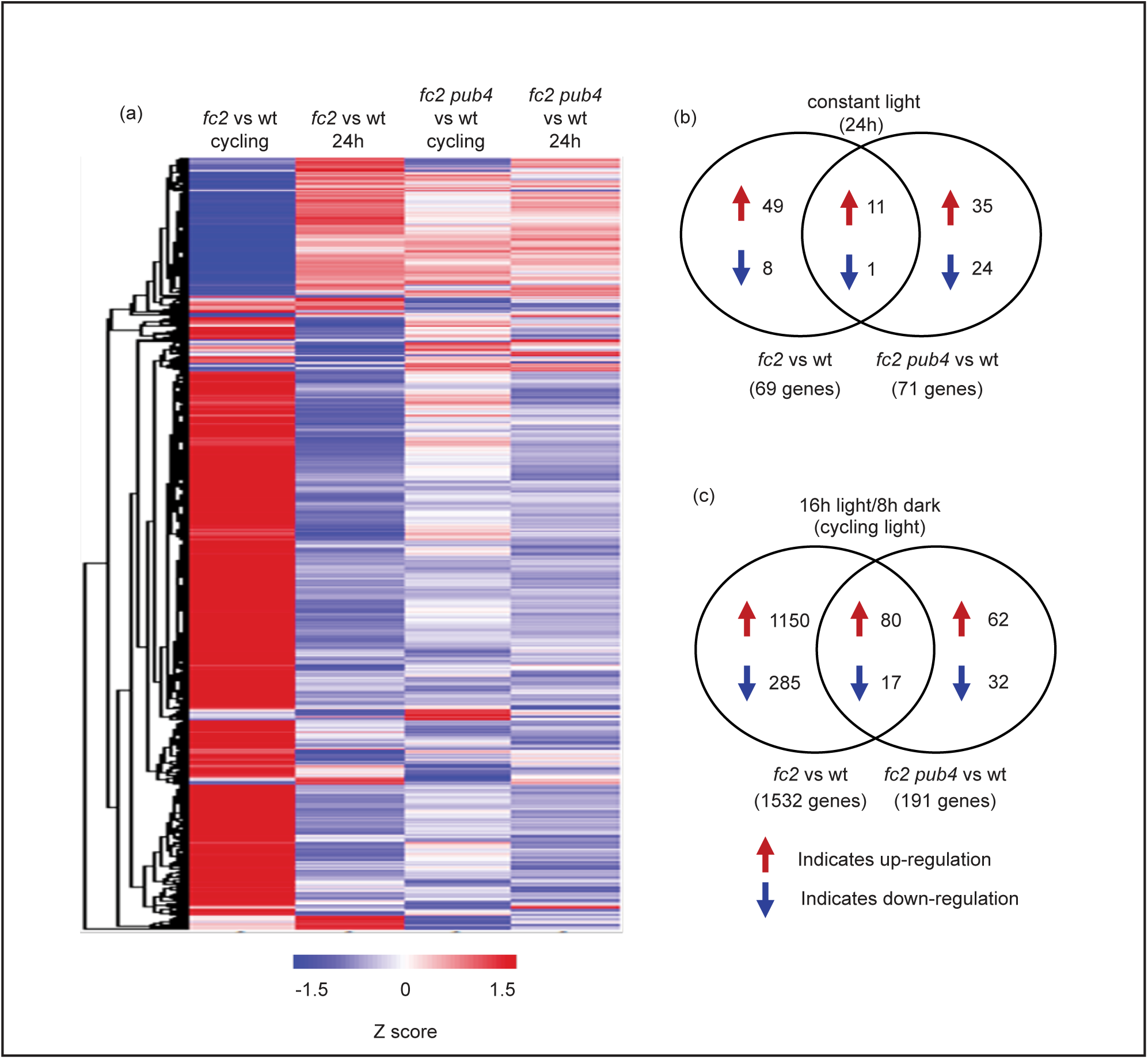
Photo-oxidative stress in *fc2* leads to a global change in transcript levels that is mostly reversed by the *pub4* mutation. An RNA-Seq analysis was performed to measure steady-state transcript levels in plants grown in permissive constant (24h) light conditions for 19 days or 17 days in constant light conditions and shifted to singlet oxygen (^1^O_2_)-producing 16h light/8h dark diurnal (cycling) light conditions for two days. (a) A heatmap representation of the 1682 differentially expressed genes (DEGs) identified by at least one pairwise comparison between genotypes (*fc2* vs. wt or *fc2 pub4* vs. wt) in one condition (24h or cycling) that passed the applied cutoff values (Log_2_FC > +/− 1.5 and a P*adj* < 0.01). The z-score color bar at the bottom shows the range of expression values, from decreased expression (blue) to increased expression (red). Genes are hierarchically clustered based on Euclidian distance of Log_2_FC and complete linkage. (b) and (c) Venn diagrams depicting unique and shared up-regulated and down-regulated DEGs in 24h light and cycling light conditions, respectively.

Under permissive 24h light conditions, only 69 genes were differentially expressed between wt and *fc2* (“*fc2* vs. wt – 24h”), with 60 (87%) showing up-regulation and 9 (13%) showing down-regulation (**Fig. 2B**). Among the up-regulated genes, there was an enrichment of gene ontology (GO) terms associated with abiotic stress (e.g., heat, response to ROS) (**Fig. S3a** and **Table S6**), but no GO-terms were enriched among down-regulated genes. Under cycling light conditions, the response in *fc2* (“*fc2* vs. wt – cycling”) was much more pronounced with 1532 genes exhibiting differential expression compared to wt. Among these, 1230 (80%) were upregulated and 302 (20%) were down-regulated (**Fig. 2c**). The up-regulated genes were enriched for GO terms broadly related to stress (predominantly biotic stress) (**Fig. 3a** and **Table S7**), including “response to jasmonic acid”, “response to salicylic acid”, “response to wounding”, and “response to bacterium”. In addition, terms related to abiotic stress such as “response to hypoxia” and “response to oxidative stress” were enriched. Conversely, the down-regulated genes were enriched for GO-terms related to photosynthesis and growth including “photosynthesis light harvesting in PSII and PSI” and “regulation of growth” (**Fig. 3b** and **Table S7**). As there are no ^1^O_2_-associated GO-terms, we compared DEGs to the list of the core ^1^O_2_ response genes (24 up-regulated and 11 down-regulated) identified by a meta-analysis previously performed by our group [22]. In *fc2* under cycling light conditions, 18 of the 24 up-regulated and 2 of the 11 down-regulated passed the cutoff values (compared to wt) (p=2.18e-18, hypergeometric test), while heatmaps show that all 35 genes followed the expected expression patterns (**Figs. 3c and d**). This expression depended on bulk ^1^O_2_ accumulation, as none of these genes were differentially expressed under 24h light conditions, compared to wt.

**Figure 3:**
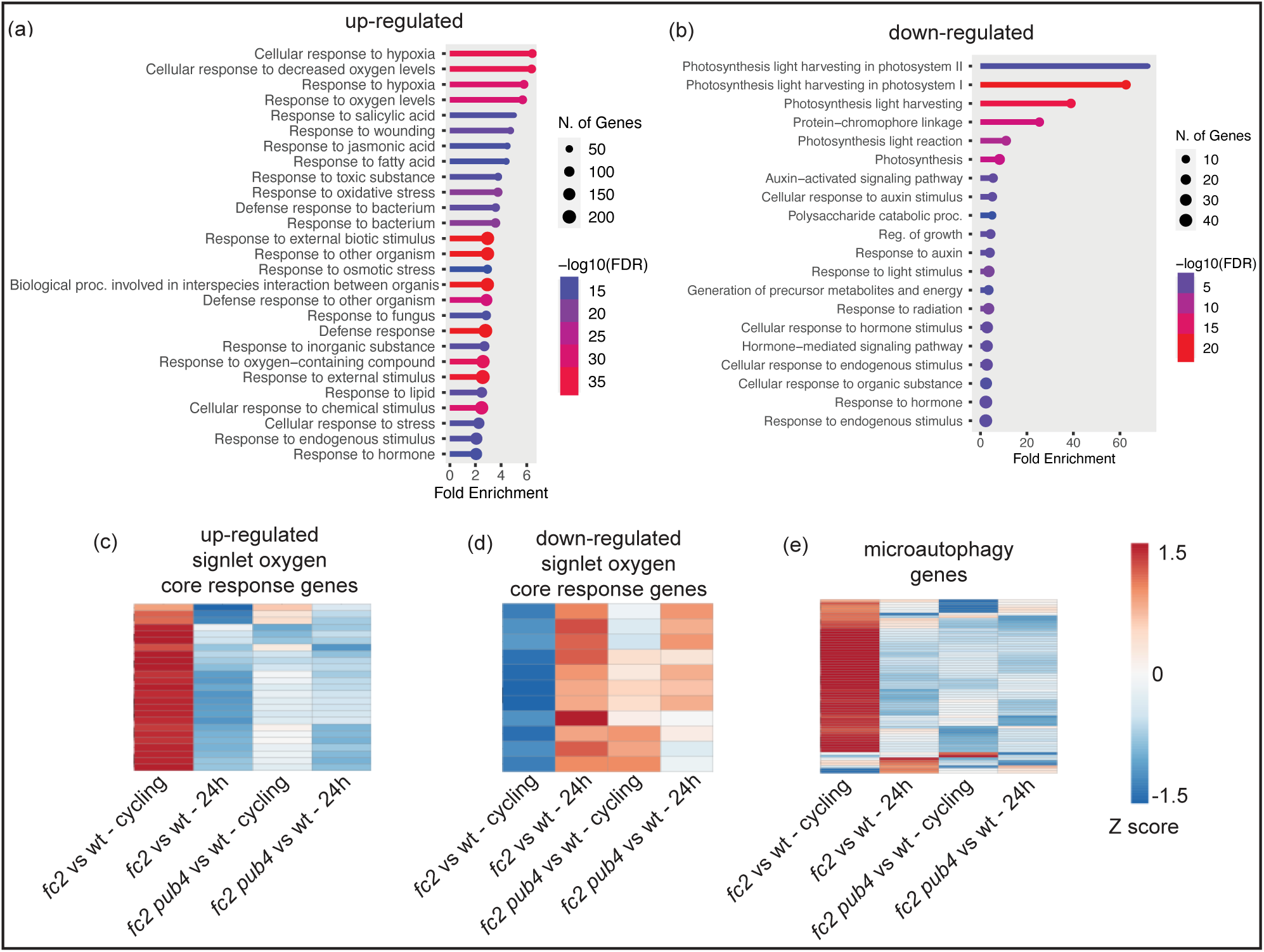
Gene ontology enrichment analysis of differentially expressed genes in *fc2*. Gene ontology (GO) term enrichment analyses of the identified (a) up- and (b) down-regulated differentially expressed genes (DEGs) in *fc2* mutants (compared to wt) in 16h light/8h dark diurnal (cycling) light conditions. Expression data is from the DESeq2 analysis of the included RNA-seq data set. GO-term enrichment analyses were performed using ShinyGO 0.80 with an FDR cutoff of p ≤ 0.01. x-axes indicate fold-enrichment. The ball size indicates the number of genes. The line colors represent FDR values. Shown are heatmaps representing the expression of (c) inducible and (d) repressible core singlet oxygen (^1^O_2_) response genes from the pairwise analyses “wt vs. *fc2*” and “wt vs. *fc2 pub4*” under cycling or constant light (24h) conditions. The z-score bar at the right shows the range of expression values, from decreased expression (blue) to increased expression (red).

Because fission-type microautophagy is expected to play a role in ^1^O_2_ induced chloroplast degradation, we monitored the expression of 66 putative markers for this degradation pathway (**Table S8**) [15]. Under 24h light conditions, 0 genes were differentially expressed in “*fc2* vs. wt – 24h” (padj ≤ 0.01) (**Fig. 3e**). However, under cycling light conditions, 38 genes were significantly induced in “*fc2* vs. wt – 24h” (p=1.27e-16, hypergeometric test), suggesting that this degradation pathway is being induced by ^1^O_2_ in photosynthetically active true leaves.

Next, we assessed the effect the *pub4* mutation has on these responses. Under 24h light conditions, 71 DEGs were identified in “*fc2 pub4* vs. wt - 24h” (46 [65%] upregulated and 25 [35%] downregulated) (**Fig. 2b**). However, only a small subset of these DEGs were shared with “*fc2* vs. wt – 24h”, indicating that the majority of the upregulated (82%) and downregulated (89%) DEGs in *fc2* are reversed by the *pub4* mutation in 24h light conditions (**Table S9**). The up-regulated genes shared (not reversed) by “*fc2* vs. wt – 24h” and “*fc2 pub4* vs. wt – 24h” were enriched for abiotic stress-related GO-terms, including “response to salt”, “response to ROS,” and “response to heat” (**Fig. S3b** and **Table S10**). Furthermore, the upregulated DEGs specific to “*fc2 pub4* vs. wt – 24h” were enriched for the “defense response” GO term (**Table S10**) suggesting that the *pub4* mutation does not entirely reverse the stress phenotype of *fc2* and may lead to a degree of constitutive stress response under permissible conditions.

In cycling light conditions, 191 DEGs were identified in “*fc2 pub4* vs. wt – cycling” with 142 (74%) showing up-regulation and 49 (26%) showing down-regulation (**Fig. 2c**). As in 24h light conditions, the majority of upregulated (93%) and downregulated (94%) DEGs found in “*fc2* vs. wt – cycling” were not shared with “*fc2 pub4* vs. wt – cycling”, (**Table S9**) suggesting that *pub4* reverses most of the gene expression in *fc2* during cycling light conditions. This was consistent for the core ^1^O_2_ response genes, none of which were differentially expressed in “*fc2 pub4* vs. wt – cycling” (**Figs. 3c and d**). *pub4* also completely reversed the induction of fission-type microautophagy genes, with 0 being induced in “*fc2 pub4* vs. wt – cycling” (**Fig. 3e**).

However, among the up-regulated DEGs shared (not reversed) by “*fc2* vs. wt – cycling” and “*fc2 pub4* vs. wt – cycling”, GO-terms associated with biotic and abiotic stress (e.g., ROS-related terms, defense responses) remained enriched (**Fig. S3c** and **Table S11**). This suggests that *fc2 pub4* plants still initiate some level of stress response, although terms associated with JA and SA signaling were notably absent. Only GO-terms associated with wax and fatty acid biosynthesis were enriched among shared down-regulated genes, while terms associated with photosynthesis were absent (**Figs. S3d** and **Table S11**). The latter observation may explain how *pub4* partially alleviates the negative impact on photosynthesis. We validated these RNA-seq data by confirming the relative expression changes of nine DEGs identified in “*fc2* vs. wt – cycling” and “*fc2 pub4* vs. wt – cycling” using RT-qPCR (**Fig. S3e**). Together, these results show the broad impact chloroplast ^1^O_2_ can have on nuclear gene expression, most of which is dependent on proper PUB4 function.

### WRKY and NAC transcription factor targets are induced by singlet oxygen and activation of NAC targets is blocked by the *pub4* mutation

Given the significant effect ^1^O_2_ has on gene expression in *fc2*, we next asked if any transcriptional regulatory mechanisms are involved. To this end, we identified additional DEGs in each of the four pairwise comparisons by relaxing our cutoff criteria (log_2_FC > +/− 1 and *padj* < 0.01) (**Table S12**) and compared the resulting DEG list to a curated list of transcription factors (TFs) (**Table S13**) [26, 27]. This led to the identification of 254 differentially expressed TFs (DETFs) in our data set (**Fig. S4a** and **Table S13**). 219 DETFs were identified in “*fc2* vs. wt – cycling”, with 216 of these DETFs possibly dependent on high levels of ^1^O_2_, as they were not found in “*fc2* vs. wt – 24h” (**Fig. S4b**). We only identified 41 DETFs in “*fc2 pub4* vs. wt – cycling”, and only 18 of these DETFs were shared with “*fc2* vs. wt – cycling”, suggesting that most of the changes observed in *fc2* can be reversed by the *pub4* mutation (**Fig. S4c**). We also analyzed these DETFs for enrichment of specific families of TFs (**Fig. S4d** and **Tables S14** and **15**). WRKY, Ethylene Response Factor (ERF), and NAC TF family member genes were significantly enriched in “*fc2* vs. wt – cycling”, while only WRKYs and ERFs were significantly enriched (albeit in reduced numbers) in “*fc2 pub4* vs. wt – cycling.” For down-regulated TFs, CONSTANS (CO)-like, basic/helix-loop-helix (bHLH), and myeloblastosis (MYB)-related TFs were enriched in “*fc2* vs. wt – cycling”, and MIKC MADS TFs were enriched in “*fc2 pub4* vs. wt – cycling.” Together these data suggest ^1^O_2_ accumulation leads to the reprogramming of hundreds of TFs. This mostly involves PUB4, although some ERFs and WRKYs may be regulated independently.

Next, we performed a GO term enrichment analysis to identify processes that may be affected by these DETFs (**Table S16**). DETFs identified in “*fc2* vs. wt – cycling” were enriched for GO-terms associated with stress hormone signaling (SA, JA, ABA, and ethylene), regulation of leaf senescence, biotic defense response (e.g., response to bacterium and chitin), and response to abiotic stresses (e.g., hypoxia and water deprivation) (**Fig. S5a**). Among down-regulated DETFs, only the GO-term “gibberellic acid signaling” was enriched. In contrast, only ethylene signaling GO-terms were observed to be enriched in up-regulated DETFs within “*fc2 pub4* vs. wt – cycling” (**Fig. S5b**) suggesting that many of the ^1^O_2_ transcriptional networks may be modulated by PUB4.

The WRKY TFs upregulated in “*fc2* vs. wt – cycling” were enriched for terms related to stress hormone signaling (SA and JA), biotic defense response (response to bacterium and chitin), response to abiotic stresses (water deprivation), and regulation of leaf senescence. The ERF TFs in the same set were enriched for terms related to ethylene stress hormone signaling, and response to abiotic stresses (water deprivation). The NAC TFs were enriched for terms associated with the regulation of leaf senescence (**Fig. S5c**). In “*fc2 pub4* vs. wt – cycling”, on the other hand, we only observed SA and ethylene stress hormone signaling to be associated with the WRKY and ERF DETFs, respectively (**Fig. S5d**). Together, these observations suggest that the *pub4* mutation negatively affects the chloroplast ^1^O_2_-induced activation of TFs involved in JA signaling, defense responses, and leaf senescence, but less so for SA signaling.

Next, we performed a transcription factor enrichment analysis (TFEA) to investigate if these induced TFs are responsible for the observed transcriptional response. We observed targets of 179 and 115 TFs in the “*fc2* vs. wt – cycling” comparison to be represented in upregulated and down-regulated DEGs, respectively (**Figs. S6a and b, Table S17**). The *pub4* mutation mostly reversed this trend with only 39 and 7 TFs shared among the up-regulated and down-regulated DEG targets, respectively, in “*fc2 pub4* vs. wt – cycling”. Among the TFs identified, WRKYs, NACs, bZIP, MYB, and BES1 TF targets were enriched in “*fc2* vs. wt – cycling” (**Fig S6c** and **Table S18**). Notably, ERF TF targets were not enriched in this analysis, even though ERF TFs were enriched in the DETF analysis. *pub4* reversed these effects, except for WRKYs, 30 of which were enriched in “*fc2 pub4* vs. wt – cycling”. Interestingly, these 30 WRKY TFs are all shared with “*fc2* vs. wt – cycling”, suggesting their function is dependent on ^1^O_2_ accumulation, but is not mediated by PUB4.

When we compared the lists of TFs from the DETF analysis and TFEA, we found an overlap of 57 and 5 TFs, respectively between “*fc2* vs. wt – cycling” and “*fc2 pub4* vs. wt cycling” (**Figs. S6d and e** and **Table S19**). This maintained the general trend of enrichment of WRKYs (23) and NACs (16) in “*fc2* vs wt – cycling”, and the retention of WRKYs (5) in *“fc2 pub4* vs. wt – cycling”. Interestingly, the five WRKYs present in “*fc2 pub4* vs. wt – cycling” that may act independently from PUB4 include WRKY33 and WRKY40, which respond to ^1^O_2_ [28–30] and SA signaling [31]. Together, these results suggest that ^1^O_2_ induces a robust transcriptional change by regulating the expression of hundreds of TFs, particularly WRKY and NAC family members. The *pub4* mutation may block part of this response by affecting the expression (WRKYs and NACs) and activation (WRKYs) of some of these TFs, while still retaining part of the PUB4-independent WRKY-directed stress response for SA signaling.

### Singlet oxygen affects plastid protein gene expression and regulation

Under cycling light conditions, downregulated DEGs in *fc2* were predominantly associated with chloroplasts, particularly the photosynthetic light reactions (**Fig. 3b**). To further understand how ^1^O_2_ affects chloroplasts and photosynthesis at the level of transcript accumulation, we used the list of 2495 plastid protein-encoding genes (PPEGs) (both plastid- and nuclear-encoded) from the Plant Proteome database (PPDB) (http://ppdb.tc.cornell.edu/dbsearch/subproteome.aspx). After removing splice variants and genes with unavailable data, a list of 1730 unique PPEGs was established and their relative expression within the RNA-seq data set were assessed (**Fig. 4a** and **Table S20**). Under cycling light conditions, 1,135 PPEGs (66%) exhibited significant (*padj* < 0.01) alterations in “*fc2* vs. to wt – cycling” (p=2.21e-266, hypergeometric test) with 975 downregulated and 160 upregulated (56% and 9% of the total PPEGs, respectively) (**Fig. 4b**). This effect was partially reversed by *pub4* with 26% of PPEGs (86 up-regulated and 361 down-regulated) still exhibiting altered expression in “*fc2 pub4* vs. wt – cycling” (p=1.07e-100, hypergeometric test).

**Figure 4:**
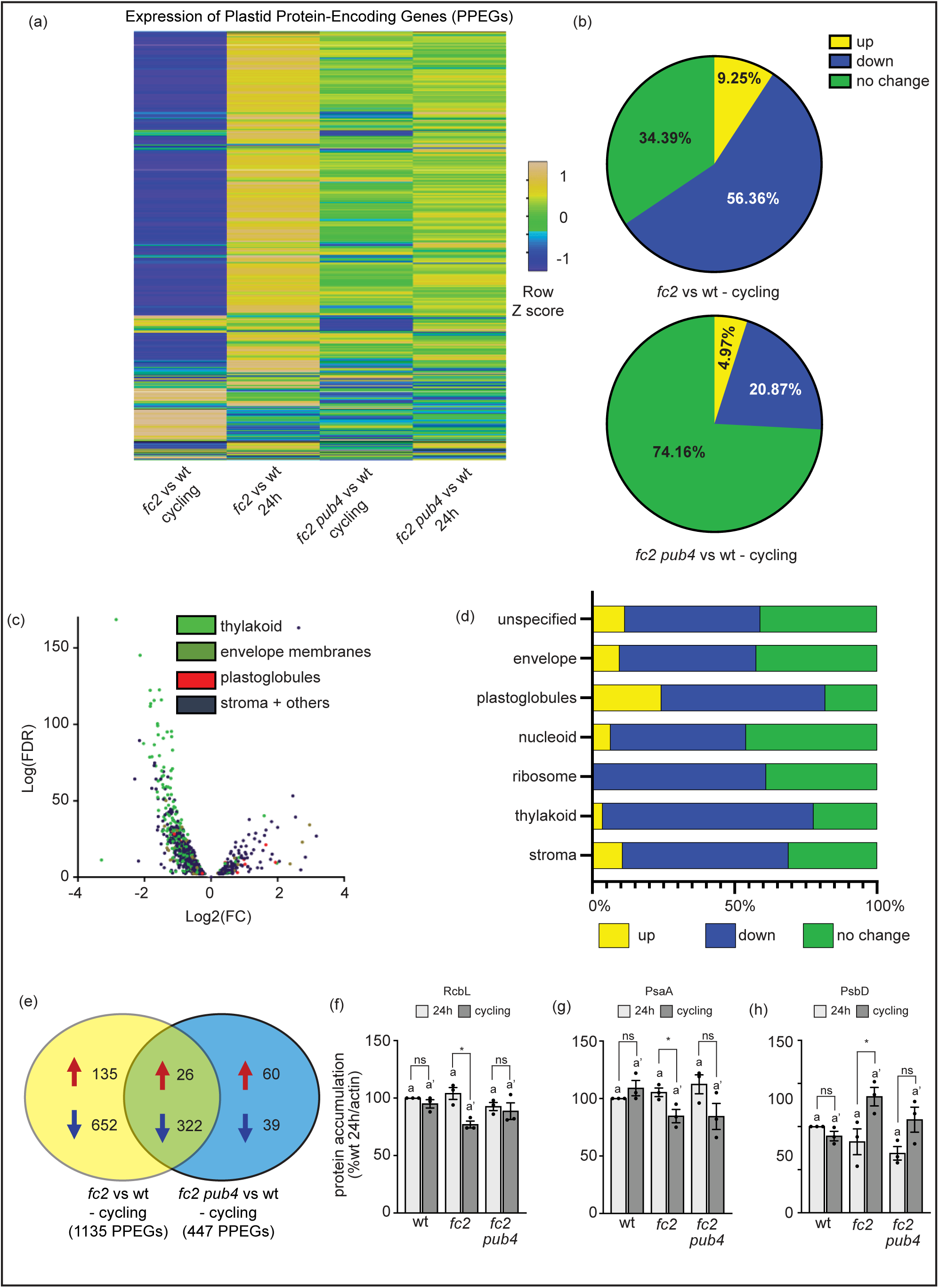
Analysis of plastid protein-encoding gene expression during singlet oxygen stress. We assessed the effect of singlet oxygen (^1^O_2_) signaling on the expression of plastid protein-encoding genes (PPEGs). Expression data is from the DESeq2 analysis of the included RNA-seq data set. (a) Heat map showing the expression of 1730 PPEGs (both plastid- and nuclear-encoded) in *fc2* and *fc2 pub4* (relative to wt), grown under constant light (24h) and 16h light/8h dark diurnal (cycling) light conditions. The up-regulated genes are shown in yellow, the down-regulated genes in blue, and genes with no change in expression are shown in green. (b) Pie chart showing the same 1730 PPEGs differentially expressed in wt vs. *fc2* (top) and wt vs. *fc2 pub4* (bottom), in cycling light conditions (*padj* < 0.01). The up-regulated genes are shown in yellow, the down-regulated genes in blue, and genes with no significant change in expression are shown in green. (c) Volcano plot representation of *PPEG* expression in *fc2* compared to wt, grown in cycling light conditions. The genes are color coded based on the predicted localization of their protein products (thylakoid, envelope membranes, plastoglobules, and stroma with other associated locations). The x- and y-axes show log_2_ fold-change in expression and the Log_10_ false discovery rate of a transcript being differentially expressed, respectively. (d) Bar graph showing percentage of PPEGs that are up-regulated (yellow), down-regulated (blue), or have no significant change (green) in *fc2* in cycling light conditions (compared to wt), based on predicted sub-plastid localization of their protein products (envelope membranes, thylakoid, stroma, plastoglobule, ribosome, nucleoid, and unspecified locations within plastid) (*padj* < 0.01). (e) Venn diagram comparing differentially expressed PPEGs in *fc2* and *fc2 pub4*, in cycling light conditions compared to wt (*padj* < 0.01). The total gene counts are grouped as up-regulated or down-regulated, indicated by a red or blue arrow, respectively. Immunoblot analysis of selected plastid-encoded proteins (f) RbcL, (g) PsaA, and (h) PsbD from leaves of three-week-old plants grown in constant (24h) light conditions or grown in 24h light conditions and shifted to 16h light/8h dark diurnal (cycling) light conditions for two days. Shown are mean values compared to actin levels (+/− SEM) and normalized to wt in 24h light conditions (n ≥ 3 leaves from separate plants). Statistical analyses were performed using one-way ANOVA tests, and the different letters above the bars indicate significant differences within data sets determined by Tukey-Kramer post-tests (P ≤ 0.05). Separate analyses were performed for the different light conditions, and the significance of the cycling light is denoted by letters with a prime symbol (ʹ). Statistical analyses of genotypes between conditions were performed by student’s t-tests (*, *P* ≤ 0.05; ns, P ≥ 0.05). Closed circles indicate individual data points.

To gain a deeper insight as to which chloroplast sub-compartments were being affected by ^1^O_2_ in *fc2* under cycling light conditions, the PPEGs were further sub-categorized by predicted protein localization (envelope, thylakoid, plastoglobule, and stroma (plus other localizations)) (**Fig. 4c**). While genes in all four categories were both up- and down-regulated in *fc2* compared to wt, genes encoding proteins in the thylakoid membrane were disproportionally down-regulated. To quantify this distribution, we further divided proteins into predicted chloroplast sub-compartments (including nucleoid, ribosome, and unspecified) and applied a significance cutoff of *padj* < 0.01 (**Fig. 4d**). As expected, the thylakoid membrane compartment is the most negatively affected in *fc2*, along with the ribosome, and nucleoid compartments. This implies that ^1^O_2_ stress in chloroplasts predominantly influences transcripts encoding photosynthetic and gene expression machinery.

To confirm this, we next performed GO-term analyses of PPEGs affected in “*fc2* vs. wt - cycling” (**Table S21**). As expected, based on protein localization, the 975 down-regulated PPEGs were enriched for GO-terms associated with photosynthesis light reactions and ribosome assembly (rRNA processing) as well as photosynthesis carbon reactions (**Fig. S7b**). The 160 up-regulated PPEGs were enriched for GO-terms associated with chorismate biosynthesis (a critical step in SA biosynthesis), senescence (chlorophyll catabolism), and amino acid synthesis (phenylalanine and tryptophan) (**Fig. S7a**). A comparison with differentially expressed PPEGs in “*fc2 pub4* vs. wt – cycling” showed that the *pub4* mutation reversed most of these patterns (**Fig. 4e**). However, both *fc2* and *fc2 pub4* were still enriched for GO-terms associated with stress (high light, cold, and heat) and photosynthesis (light and carbon reactions) among up- and down-regulated DEGs, respectively (**Fig. S7c and d**). Thus, the *pub4* mutation lessens the negative impact on photosynthesis even though many of the same categories of stress genes are still induced. This is consistent with *fc2 pub4* plants remaining greener and experiencing less chlorophyll cellular degradation compared to *fc2* (**Figs. 1a-d**).

Some differentially expressed PPEGs in “*fc2 pub4* vs. wt – cycling” were unique. Among the 60 up-regulated PPEGs, there was enrichment for GO-terms related to plastid gene expression (transcription, translation, rRNA processing), nucleotide import, and protein import (**Fig. S7e**). Conversely, the 39 down-regulated PPEGs were enriched for GO terms related to SA biosynthesis (i.e., chorismate biosynthesis) and aromatic amino acid synthesis (**Fig. S7f**). These latter terms were associated with the upregulated PPEGs in “*fc2* vs. wt – cycling” and suggest that the *pub4* mutation leads to an increase in plastid gene expression and chloroplast development while down-regulating SA biosynthesis genes.

To test if PPEGs in the nucleus or plastid are more affected by ^1^O_2_ stress conditions, we compared expression patterns of plastid (40 genes) and nuclear genes (51) encoding components of the photosynthetic light reactions (as well as *rbcL* in the plastid) **(Figs. S8a and b)**. Of the plastid genes, only three genes encoding ATP synthase subunits were differentially expressed in “*fc2* vs. wt – cycling” (*padj* < 0.01) (p=0.131, hypergeometric test). In contrast, all 51 nuclear genes were repressed in “*fc2* vs wt - cycling” (p=2.08e-46, hypergeometric test). For each of these genes, the *pub4* mutation reduced the strength of the repression, but 44 were still significantly repressed in “*fc2 pub4* vs. wt - cycling” (p=2.05e-49, hypergeometric test).

Next, we tested if plastid-encoded gene expression correlated with protein abundance. Immunoblotting was used to compare the accumulation pattern of the plastid-encoded proteins RbcL (large subunit of RuBisCO), PsaA (PSI complex subunit), PsbD/D2 (PSII complex subunit), and PetB (cytochrome b_6_f complex subunit). Despite a mild effect on plastid-encoded transcripts, both RbcL and PsaA were less abundant in *fc2* plants in cycling light, compared to 24h light conditions (**Figs. 4f and g**). PetB accumulation, on the other hand, was not significantly altered (**Figs. 8f**). Surprisingly, the D2 protein was more abundant in cycling light-stressed plants (**Fig. 4h**). These effects were slightly reversed by *pub4*, although the D2 protein still accumulated to higher levels in *fc2 pub4* plants grown under cycling light conditions. We also tested protein accumulation in plants after seven days in cycling light conditions and observed similar patterns (**Figs. S8c-e, g**). Together, this suggests that ^1^O_2_ affects nuclear and plastid-encoded transcripts differently, and may also affect plastid-encoded protein levels through post-translational regulatory mechanisms.

### Singlet oxygen stress in the *fc2* mutant leads to the induction of salicylic acid synthesis and signaling

The GO-terms “response to SA” and “chorismate biosynthesis” were strongly enriched among genes and TFs upregulated in *fc2* in response to ^1^O_2_ stress (**Figs. 3a, S5a and c**, **S7a** and **Table S7**) leading us to hypothesize that SA levels and/or signaling may also be induced. To this end, we assessed the relative expression of ten genes associated with SA synthesis (**Figs. 5a and b** and **Table S22**) [32]. In “*fc2* vs. wt – 24h,” none of these genes were significantly induced (*padj* < 0.01). Under cycling light conditions, however, four of these genes were significantly induced in “*fc2* vs. wt - cycling.” This included *ISOCHORISMATE SYNTHASE1* (*ICS1*) and *PBS3*, but none of the genes encoding phenylalanine ammonia-lyase (*PAL1-4*). Thus, it appears the dominant ICS pathway, rather than the PAL pathway (**Fig. 5a**), is induced in *fc2* under cycling light conditions. This response was mostly reversed by the *pub4* mutation. Some of the ICS pathway genes were mildly upregulated in “*fc2 pub4* vs. wt – 24h/cycling” but these differences were not statistically significant.

**Figure 5:**
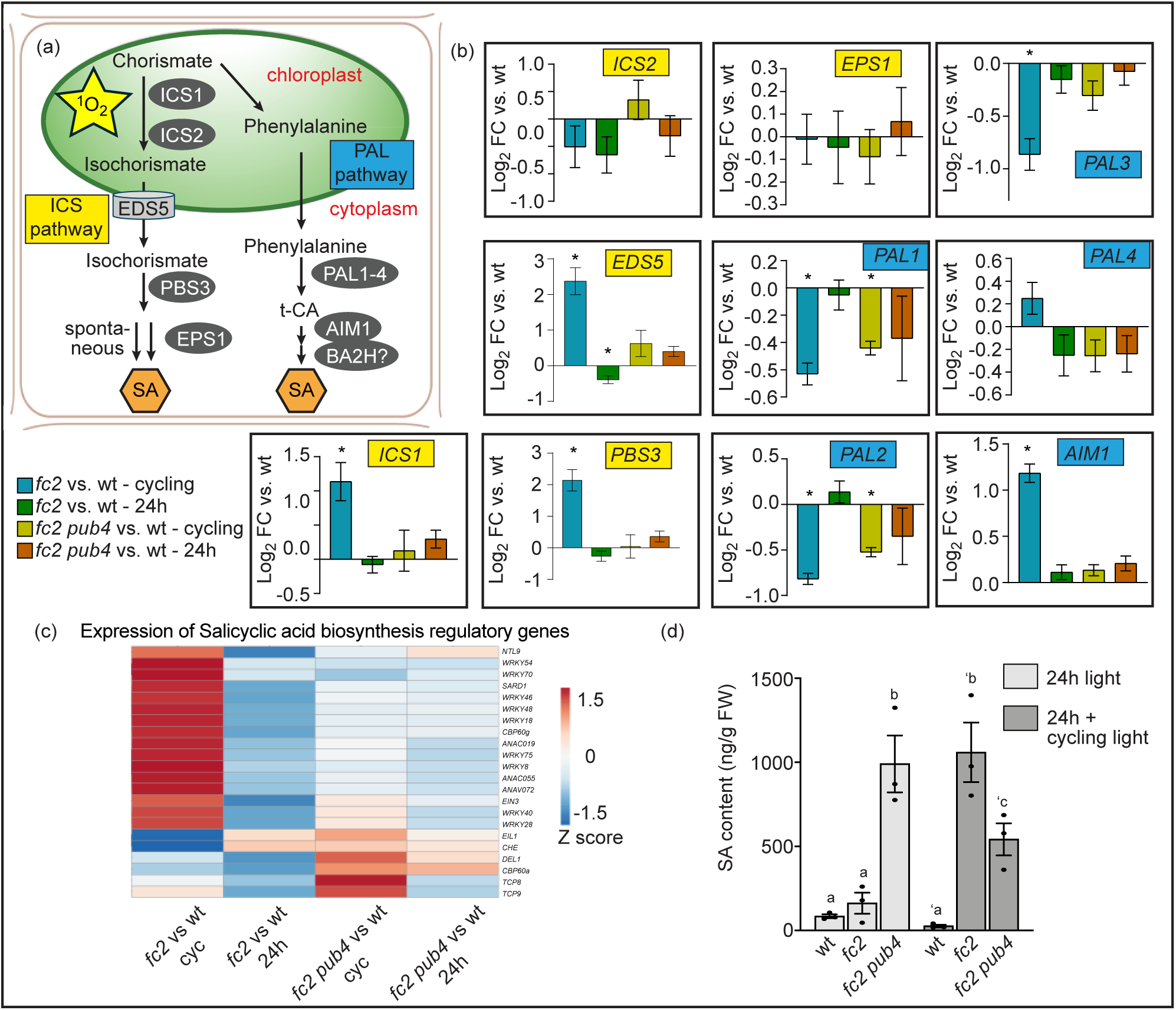
Singlet oxygen and *pub4* alter the expression of salicylic acid-related genes and salicylic acid content. The effect of singlet oxygen (^1^O_2_) on salicylic acid (SA) content and signaling was assessed. (a) An overview of SA synthesis via the isochorismate synthesis (ICS) (left side) and the phenylalanine ammonia-lyase (PAL) pathway (right side). (b) Relative expression (compared to wt in the same light condition) of ten key SA biosynthesis genes from panel a. Genes involved in the ICS and PAL pathways are labeled in yellow and blue, respectively. Expression data is from the DESeq2 analysis of the included RNA-seq data set. Error bars in the graph represent standard error (ifcSE) estimated by DESeq2 (*, *padj* < 0.01). (c) Heatmap showing relative expression of genes associated with the regulation of SA biosynthesis from the RNA-seq data set (“wt. vs. *fc2*” or “wt vs. *fc2 pub4*” in constant 24h light conditions or after two days of 16h light/8h dark diurnal (cycling) light conditions). The blue and red colors correspond to low and high gene expression, respectively. (d) SA content (ng/g fresh weight (FW)) measured in 17-day-old plants (grown in 24h light conditions or after two days of cycling light conditions). Shown are mean values ± SEM (n = 3 replicates). Statistical analysis was performed by one-way ANOVA followed by Tukey’s multiple comparison test. Different letters above the bars indicate significant differences (*P* value ≤ 0.05). Closed circles represent individual data points.

We also measured the expression of 22 genes associated with the regulation of SA biosynthesis [33] and observed a similar pattern (**Fig. 5c** and **Table S22**). In constant light conditions, the expression of only one gene in *fc2* was significantly altered (“*fc2* vs. wt – 24h”, *padj* < 0.01, p=0.248, hypergeometric test). However, under cycling light conditions, 17 genes (77%) were significantly affected (15 up-regulated and 2 down-regulated) in “*fc2* vs. wt – cycling” (p=3.29e-11, hypergeometric test). This effect was not strongly reversed by *pub4*, with 6 (WRKYs 8, 28, 20, and 49) and 2 (WRKYs 28 and 46) of these genes were still differentially expressed in “*fc2 pub4* vs. wt – cycling” and “*fc2 pub4* vs. wt – 24h,” respectively (p=3.89e-4, hypergeometric test). This indicates that *pub4* may not completely block SA regulation by ^1^O_2_, which aligns with the TF analysis (**Fig. S5d**).

Next, we measured SA levels in 17-day-old plants to determine if these changes in transcript levels led to increased hormone levels. Under constant light conditions, there was no significant difference in SA content between *fc2* and wt (**Fig. 5d**). However, upon exposure to cycling light conditions for two days, *fc2* accumulated ∼40-fold more SA content than wt. In line with the transcript analysis, the *pub4* mutation reduced this induction by 50% but *fc2 pub4* mutants still accumulated significantly more SA than wt. Furthermore, SA content increased more than 20 times in *fc2 pub4* under constant light conditions. This suggests that SA content is constitutively high in *fc2 pub4* plants despite only a mild effect on some genes involved in SA synthesis (**Fig. 5b**) and regulation (**Fig. 5c**). To test if the high level of SA in *fc2 pub4* plants was leading to a canonical SA response, we identified DEGs between *fc2* and *fc2 pub4* in constant light conditions (*padj* < 0.01) and compared them to 204 SA-responsive genes (**Table S23)** [34]. 48 (24%) of these genes were differentially expressed between *fc2* and *fc2 pub4* (42 [88%] and 6 [12%] upregulated and downregulated, respectively) (p=4.21e-28, hypergeometric test). Thus, the high level of SA in *fc2 pub4* is activating a mild, but constitutive transcriptional SA response.

### Singlet oxygen stress leads to the induction of jasmonic acid synthesis and signaling

^1^O_2_-stressed *fc2* mutants showed an enrichment of genes associated with jasmonic acid (JA) synthesis and metabolism (**Fig. 3a**), suggesting that JA synthesis may also be induced by ^1^O_2_ accumulation. To test this, we used the RNA-seq data set to analyze the expression of 11 key genes involved in JA biosynthesis and signaling (**Fig. 6a** and **Table S24)**. Eight of these genes (including seven involved in biosynthesis) were significantly upregulated in “*fc2* vs. wt – cycling,” and only *DAD1* was still induced in “*fc2 pub4* vs. wt – cycling” (*padj* < 0.01). To test if this change in gene expression leads to altered JA levels, plants were assessed for JA content (**Fig. 6b**). Under 24h conditions, all genotypes accumulated similar levels of JA. After being shifted to cycling light conditions for two days, *fc2* plants accumulated ∼6-fold more JA than wt. The *pub4* mutation reversed this phenomenon.

**Figure 6:**
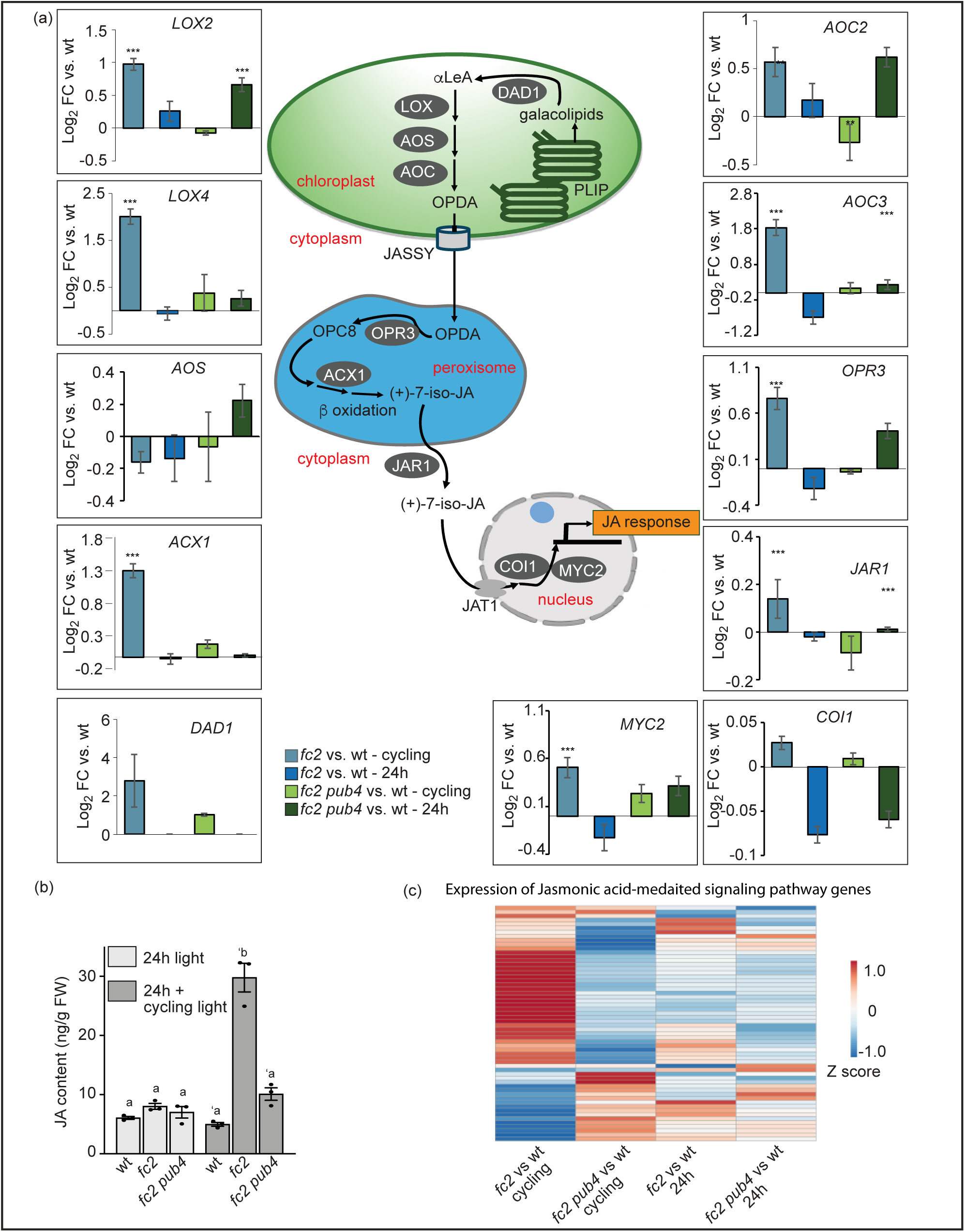
Jasmonic acid biosynthesis and signaling are induced in *fc2* under cycling light conditions. The impact of singlet oxygen (^1^O_2_) on jasmonic acid (JA) biosynthesis and signaling was assessed in constant light (24h) and 16h light/8h dark diurnal (cycling) light conditions. (a) Schematic illustration of the JA biosynthesis pathway, with relative expression (compared to wt in the same light condition) of 11 key JA biosynthesis and signaling genes. Expression data is from the DESeq2 analysis of the included RNA-seq data set. Error bars in the graph represent standard error (ifcSE) estimated by DESeq2 (**, *padj* < 0.01; ***, *padj* < 0.001). In chloroplasts, Defective In Anther Dehiscence 1 (DAD1) catalyzes the conversion of galactolipids into α-linolenic acid (α-LeA), which is converted into 12-oxo-phytodienoic acid (OPDA) by a series of enzymes; 13-Lipoxygenase (LOX), Allene Oxide Synthase (AOS), and Allene Oxide Cyclase (AOC). Subsequently, OPDA is exported out of the chloroplast via the channel protein JASSY. Inside the peroxisome, OPDA is reduced by the OPDA reductase 3 (OPR3) and shortened in the carboxylic acid side chain by β-oxidation enzymes (ACX1) into JA. In the cytosol, Jasmonate Resistant 1 (JAR1) conjugates isoleucine to JA converting it into JA-Ile. Jasmonate Transporter 1 (JAT1) transports JA-Ile into the nucleus. In response to stressed conditions, JA-Ile attach with the COI1-SCF (coronatine insensitive1-Skp1-Cul1-F-box), to promote ubiquitination and degradation of jasmonic acid repressors JAZ and JAV1, thereby activating JA response genes. (b) Graph showing JA content (ng/g fresh weight (FW)) measured in whole rosettes from plants grown for 19 days constant (24h) light conditions or 17 days in 24h light conditions and two days of 16h light/8h dark diurnal (cycling) light conditions. Values are means ± SEM (n = 3 biological replicates). Statistical analyses in b were performed using one-way ANOVA tests, and the different letters above the bars indicate significant differences within data sets determined by Tukey-Kramer post-tests (P ≤ 0.05). Separate analyses were performed for the different light treatments, and the significance of cycling light treatment is denoted by letters with a prime symbol (ʹ). (c) Heatmap showing relative expression of 58 genes from gene ontology term “JA mediated signaling pathway” (GO:0009867). The blue and red colors correspond to low and high gene expression, respectively.

Next, we used the RNA-seq data set to analyze expression of 58 genes associated with the GO term “JA mediated signaling pathway” (GO:0009867) ((**Fig. 6c** and **Table. S22**). Only one gene was differentially expressed (*padj* < 0.01) in “*fc2* vs. wt – 24h” (p=0.370, hypergeometric test). However, under cycling light conditions, 37 genes were differentially expressed (23 up and 14 down) in “*fc2* vs. wt - cycling” (p=2.81e-9, hypergeometric test). This included *MYC2,* a “master switch” of JA signaling [35, 36]. Furthermore, the *pub4* mutation reversed this response, with only 8 of these selected JA genes in “*fc2 pub4* vs. wt – cycling” being differentially expressed (p=0.076, hypergeometric test). Thus, the upregulation of JA biosynthesis genes, elevated levels of JA content, and differential expression of signaling genes indicate that JA signaling is triggered in response to ^1^O_2_ stress in *fc2* mutants and blocked by the *pub4* mutation.

### Senescence is induced in singlet oxygen-stressed *fc2* plants

When *fc2* mutants are subjected to cycling light conditions, they develop senescence-like phenotypes, characterized by pale leaves, an increase in plastoglobules, degradation of chloroplasts, and cell death. These phenotypes are reversed by the *pub4* mutation [12, 14, 15]. We also observed an enrichment of senescence-related GO-terms in genes up-regulated in “*fc2* vs. wt - cycling” (but not in “*fc2 pub4* vs. wt - cycling”) (**Tables S7** and **S11**). To investigate if canonical senescence pathways are activated by ^1^O_2_ signals, we measured the expression of genes involved in regulating leaf senescence within the RNA-Seq data set. We compared expression patterns of 128 genes, associated with the GO-term “leaf senescence” (GO: 0010150) (**Fig. S9a** and **Table S25**). Only 2% were up-regulated in “*fc2* vs. wt - 24h” (p=0.249, hypergeometric test), but 47% were up-regulated in “*fc2* vs. wt – cycling” (*padj* < 0.01) (p=2.19e-19, hypergeometric test). This effect in cycling light conditions was mostly reversed by the *pub4* mutation with only 11% of genes being induced in “*fc2 pub4* vs. to wt – cycling” (p=3.26e-4, hypergeometric test).

To refine our analysis, we selected 38 senescence-associated and regulatory genes from the literature (**Fig. 7a** and **Table S25**) [37, 38]. In 24h light conditions, none of these genes were differentially expressed in “*fc2* vs. wt – 24h” (*padj* < 0.01) (p=0.548, hypergeometric test). However, under cycling light conditions, most of these genes were induced (24) or repressed (4) in “*fc2* vs. wt - cycling” (p=1.78e-9, hypergeometric test). Two of the repressed genes, *CAB1* and *RBCS1A*, are also repressed during senescence [37]. Notably, four of the five *SENESCENCE-ASSOCIATED GENES* (*SAGs*) tested were induced in “*fc2* vs. wt – cycling.” The significant up-regulation of genes encoding the TFs (and positive regulators of senescence [39, 40]) *ANAC016*, *ANAC019*, *ANAC72*, and *WRKY53* in “*fc2* vs. wt – cycling,” suggests that they may be initial targets of the ^1^O_2_ signal. Additionally, *POLYAMINE OXIDASE 1* (*PAO1)*, encoding an enzyme involved in chlorophyll catabolism [41], was induced. These patterns were reversed by the *pub4* mutation (only 5 and 1 were induced or repressed, respectively, in “*fc2 pub4* vs. wt – cycling”) (p=0.076, hypergeometric test). Together, these data suggest that senescence pathways are induced in *fc2* mutants under ^1^O_2_ accumulating conditions. This was reversed by *pub4*, possibly due to a decrease in JA levels and/or by blocking the initiation of senescence pathways.

**Figure 7:**
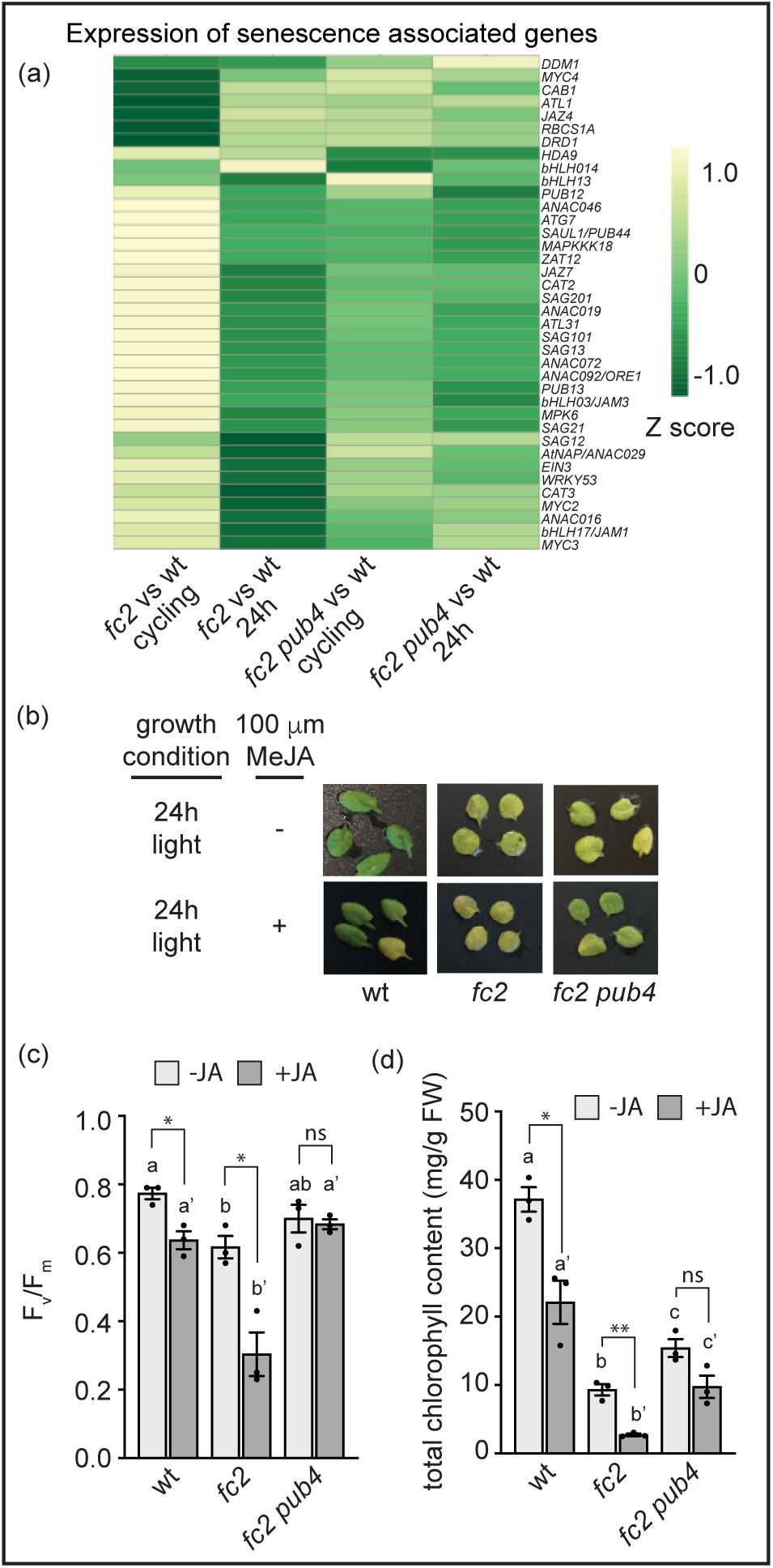
Activation of senescence pathways is blocked by the *pub4* mutation. Senescence is induced in *fc2* mutants under cycling light conditions and blocked by the *pub4* mutation. (a) Expression data from the DESeq2 analysis of the included RNA-seq data set was used to create a heatmap of genes associated with senescence. Increased and decreased expression relative to wt is indicated by yellow and green, respectively. (b) Representative images of detached 3rd and 4th rosette leaves from plants grown in constant (24h) light conditions. The leaves were incubated in the dark with water (control conditions) or with 100 μM methyl jasmonate (MeJA) for 3 days. (c) Measurement of the maximum quantum efficiency of photosystem II (F_v_/F_m_) of leaves in panel b. (d) Total chlorophyll content of leaves in panel b. Statistical analyses were performed using one-way ANOVA tests, and the different letters above the bars indicate significant differences within data sets determined by Tukey-Kramer post-tests (P ≤ 0.05). Separate analyses were performed for the different treatments, and the significance of the MeJA treatment is denoted by letters with a prime symbol (ʹ). Statistical analyses of genotypes between conditions were performed by student’s t-tests (*, *P* ≤ 0.05; **, P ≤ 0.01; ns, P ≥ 0.05). n = 3 whole leaves from separate plants. Error bars = +/− SEM. Closed circles indicate individual data points.

To test this assertion, we induced early leaf senescence in the absence of photo-oxidative stress by incubating detached leaves in the dark for three days [42]. If plants were first grown in 24h light conditions, leaves from all three genotypes exhibited some level of senescence as evidenced by the visual yellowing of leaves (**Fig. 7b**). This pattern did not significantly change if plants were initially gown under 24h light and shifted to cycling light conditions (**Fig. S9b**). Next, we measured the maximum photosynthetic efficiency of PSII (F_v_/F_m_) values of these leaves. Without dark incubation, F_v_/F_m_ values were similar in all genotypes (**Fig. S9c**). After dark incubation, the values decreased, but to different levels. F_v_/F_m_ values were highest for wt (mean = 0.77) and lowest for *fc2* mutants (mean = 0.61), while *fc2 pub4* was in between (mean = 0.7) (**Fig. 7c**). Total chlorophyll content was significantly reduced in *fc2* compared to wt, while the *pub4* mutation significantly reversed this trend (**Fig. 7d**). A similar pattern was observed with leaves from plants exposed to cycling light conditions (**Figs. S9d and e**). Together, these results suggest *fc2* mutants are sensitive to dark-induced senescence, which is delayed by *pub4*.

To enhance senescence further, we added 100 μM methyl jasmonate (MeJA) before dark incubation [42]. Under these conditions, both wt and *fc2* leaves showed pronounced yellowing, whereas *fc2 pub4* leaves showed less visual senescence (**Fig. 7b**). For wt and *fc2*, F_v_/F_m_ values and total chlorophyll content were significantly reduced by JA, and both were significantly lower in *fc2* compared to wt (**Figs. 7c and d**). *pub4* not only reversed this trend (*fc2 pub4* had higher F_v_/F_m_ values and total chlorophyll content compared to *fc2*), but JA did not significantly reduce these values in the *fc2 pub4* mutant. Similar results were obtained with plants exposed to cycling light conditions (**Figs. S9d and e**). Together, these results suggest that *pub4* delays the induction of senescence pathways whether they are activated by photo-oxidative damage or dark/JA treatments.

## Discussion

### Singlet oxygen specifically impacts chloroplast function

A large body of evidence has demonstrated that chloroplast ^1^O_2_ in *fc2* mutants can trigger multiple cellular signals affecting gene expression, chloroplast degradation, and PCD [6–8], but these signals are poorly understood. Here, we demonstrate that the accumulation of ^1^O_2_ in mature *fc2* chloroplasts mutants leads to a robust retrograde signal significantly affecting RNA profiles (1,532 DEGs in *fc2* vs wt) (**Figs. 2a and c**). There was a clear impact on chloroplast function as the repressed genes (302 DEGs) were strongly enriched for photosynthesis (**Fig. 3b**). In addition, more than half of plastid protein-encoding transcripts (1,135 of 1,730) were also repressed in *fc2* compared to wt (**Fig. 4b**). This effect on the transcriptome was partly dependent on proper PUB4 function (**Figs. 2a and 4b**). Due to the cytoplasmic localization of PUB4 [12, 43], this observation further supports the existence of a secondary ^1^O_2_ signal exiting the chloroplast to regulate chloroplast function accordingly, possibly by mitigating further production of ROS or by redirecting resources. Interestingly, PPEGs involved in chloroplast development (e.g., plastid gene expression, plastid protein import) were upregulated in *fc2 pub4* mutants in cycling light conditions (**Fig. S7d**), suggesting that these chloroplasts may have increased protein turnover and synthesis that could offer protection from photo-oxidative damage. Chloroplast ^1^O_2_ accumulation due to a far-red block of chloroplast development was also reported to repress nuclear-encoded photosynthesis genes [44], further supporting a signaling role for ^1^O_2_ to limit photosynthesis. However, the far-red block signal was dependent on *EXECUTER1* (*EX1*) and *EX2*, which are dispensable in adult *fc2* plants [12, 22]. These findings, along with the only partial reversion of *PPEG* expression by the *pub4* mutation (**Fig. 4e**), support a model where multiple ^1^O_2_ signals impact photosynthetic gene expression via retrograde signaling.

### Transcription factor networks are involved in propagating the singlet oxygen signal

The large change in ^1^O_2_-induced gene expression included the induction of WRKY, ERF, and NAC TF families (**Figs. S4a and b**), consistent with stress responses involving SA and JA [45] and the induction of leaf senescence [46]. A complementary TFEA analysis was used to determine which transcription factors were responsible for the observed DEGs and confirmed that the WRKY and NAC TF families play important roles in propagating the ^1^O_2_ signal (**Fig. S6c**). However, ERF targets were not found to be enriched, suggesting that this class of TFs may not be as active during ^1^O_2_ signaling.

The TFEA also revealed some insight into the role of PUB4 in signaling. Both NAC TF transcripts and NAC family target transcripts are reversed by *pub4*, indicating that such NAC TF networks may regulate ^1^O_2_-induced PCD or senescence (**Fig. S6c**). However, the induction of WRKY targets is only modestly affected by *pub4*, despite the reversal of WRKY TF transcript accumulation. This suggests that ^1^O_2_ affects at least some WRKY function independently from PUB4. One of the WRKYs still active in *fc2 pub4* is WRKY33, which has been reported to play a role in defense against necrotrophic pathogens by regulating crosstalk between SA- and JA-response pathways [47, 48] and is an early ^1^O_2_ response gene [29]. Another TF, WRKY40, may be involved in the EX1-dependent regulation of ^1^O_2_-responsive genes [30]. Thus, some PUB4-independent ^1^O_2_ signaling occurs through WRKYs, but it does not trigger PCD. Such signaling could possibly occur through the production of β-cyclocitral (β-cc), a β-carotene oxidation product, that can induce a ^1^O_2_ retrograde signal, but does not induce PCD [49]. Thus, ^1^O_2_ may initiate the induction of parallel signals involving distinct sets of TFs, with only part of this signal being modulated by PUB4 and leading to PCD and senescence.

### Singlet oxygen induces stress hormone synthesis, which is altered by the *pub4* mutation

In *fc2* plants, ^1^O_2_ stress led to an induction of genes and TFs involved in SA and JA biosynthesis (**Figs. 3a and S5a and c**) as well as a concomitant increase in SA and JA levels (**Figs. 5d and 6b**). The *pub4* mutation blocked the accumulation of JA, but SA levels remained elevated in *fc2 pub4* mutants. Moreover, *fc2 pub4* mutants were observed to have constitutively elevated SA levels and signaling (**Fig. 5d**) under permissive 24h light conditions. Together, these results indicate that PUB4 plays a role in modulating both hormones and their downstream responses. This is significant as previous studies have linked both SA and JA as playing important roles in abiotic stress signaling, particularly in regulating PCD in response to photosynthetic stress [7]. EL stress, which can lead to the production of both ^1^O_2_ and hydrogen peroxide in chloroplasts, leads to the induction of SA and JA synthesis and signaling [50, 51]. As EL is a complex stress that often includes heat stress, it is not known which ROS in which cellular compartment may be inducing these responses, although chloroplast ^1^O_2_ accumulation in the conditional *flu* mutant has also been linked to increased JA and SA levels [52, 53]. The impact of these hormones on PCD has been reported, but with conflicting results; different reports suggesting that these hormones can either inhibit or induce PCD [52, 54]. Thus, it is possible that PUB4 can affect ^1^O_2_ signaling via modulation of SA and/or JA signaling, but additional studies will be needed to understand these connections.

It is not clear why *pub4* mutants accumulate high levels of SA. Transcripts of SA biosynthesis genes are only modestly increased in *fc2 pub4*, so it is tempting to speculate that the E3 ligase activity of PUB4 targets an enzyme involved in SA metabolism and regulates this pathway at a post-translational level. However, no in vivo PUB4 ubiquitination targets have been identified (in vitro, PUB4 has been shown to ubiquitinate BIK1, but this activity is not required for its function in innate immunity [55, 56]). This is surprising considering PUB4 has been shown to have autoubiquitination activity in vitro [43] and is involved in multiple (and seemingly incompatible) cellular processes including stress signaling [12, 17], pollen development [43], meristem development [57], and pattern triggered immunity [55, 56]. Some of this complexity of mutant phenotypes may be due to different *pub4* mutant alleles being used in these studies. Here, we have used *pub4-6*, a semi-dominant point mutation predicted to affect the U-Box motif produce a protein unable to interact with its E2 substrate [12]. Null T-DNA *pub4* alleles, which are used in studies connecting PUB4 function to development and immunity [43, 55–57], are unable to block ^1^O_2_ signaling [12] suggesting that the protein produced by the *pub4-6* allele likely has a unique effect on stress physiology. Nonetheless, a better understanding of the molecular function of PUB4 should shed light on its ability to affect SA levels and how this protein is connected to so many points of cellular physiology.

### The *pub4* mutation blocks senescence and cellular degradation pathways

In *fc2* plants, ^1^O_2_ stress also led to an induction of genes and TFs involved in senescence (**Figs. 7a and S5a and c**) and fission-type microautophagy (**Fig. 3e**), which were strongly blocked by the *pub4* mutation, again supporting the common model that ^1^O_2_-induced cellular degradation is the result of a genetic signal [6–8]. Because *pub4* also reduced JA accumulation, we hypothesized that *pub4* may generally block senescence and cellular remobilization pathways. Indeed, when senescence was induced by a combination of carbon starvation and JA treatment, the *pub4* mutation led to a “stay green” phenotype with delayed senescence (**Fig. 7b-c**). Thus, PUB4 may also act downstream of JA accumulation during stress to promote senescence and cellular breakdown. This observation is somewhat contradicted by the early senescence phenotype of *pub4* mutants under permissive light conditions [12, 23]. However, similar to autophagy (*atg*) mutants [58], this senescence (which is activated after 30 days in *pub4* and is later than timepoints used in this study) may be a consequence of impaired cellular turnover machinery as a seedling. How the *pub4* mutation affects such degradation pathways is not clear, but is likely connected to its impact on retrograde signaling and gene expression in response to photo-oxidative stress. However, as an E3 ligase, at least part of PUB4’s impact on stress signaling is also expected to involve post-translational regulation. How this activity is related to gene expression and/or cellular degradation in response to stress should be the focus of further studies.

## Conclusions

In the conditional ^1^O_2_ accumulating mutant, *fc2*, ^1^O_2_ leads to rapid cellular degradation and PCD, which is dependent on normal PUB4 function. Here we show this ^1^O_2_ also leads to a global change in transcript accumulation affecting photosynthesis, chloroplast function, stress hormone (SA and JA) synthesis, fission-type microautophagy, and senescence, involving a transcription factor network enriched with WRKY and NAC TFs. SA and JA levels are also highly elevated and are predicted to play a role in these processes. Most of these effects are reversed by the *pub4* mutation, suggested that at least part of the *pub4* phenotype is due to blocking ^1^O_2_ retrograde signals to the nucleus. However, we also demonstrate that *pub4* mutants have constitutively high levels of SA and block JA-induced senescence pathways, suggesting that PUB4 likely plays a complex role in regulating stress signaling and PCD in response to photo-oxidative stress.

## Methods

### Plant material and growth conditions

The *Arabidopsis thaliana* ecotype Columbia-0 was used as wild type (wt). Supplementary **Table S26** lists the mutant lines used. Mutant line *fc2-1* (GABI_766H08) [59] came from the GABI-Kat [60] collection and *fc2-1 pub4-6* was previously described [12]. **Table S27** lists the primers used.

Seeds were sterilized using 30% liquid bleach (v/v) with 0.04% Triton X-100 (v/v) for 5 min, washed three times with sterile water, and resuspended in a 0.1% agar solution. Resuspended seeds were spread on plates containing Linsmaier and Skoog medium pH 5.7 (Caisson Laboratories, North Logan, UT, USA) with 0.6% micropropagation type-1 agar powder. Following 3–5-day stratification at 4°C in the dark, seeds were germinated in either 24h constant light (permissive) conditions or 6h light / 18h dark cycling light (stress conditions) at 120 μmol photons m^-2^ sec^-1^ at 21°C in fluorescent light chambers (Percival model CU-36L5).

For adult plant experiments, seven-day-old plants grown in 24h constant light conditions were transferred to PRO-MIX LP15 soil and growth was continued under the same growth conditions in a fluorescent reach-in plant growth chamber (Conviron). To induce ^1^O_2_ stress, adult plants were shifted to cycling light conditions (16h light / 8h dark).

### RNA-Seq analyses

For whole transcriptome RNA-Seq analyses, plants were grown under 24h constant light conditions for 16 days. One set of plants was then shifted to cycling light conditions (16h light / 8h dark) for two days. Total leaf tissue was collected from 18-day-old plants one hour after subjective dawn. Samples were collected in biological triplicates, each replicate consisting of leaves pooled from three plants. Total RNA was extracted using the RNeasy Plant Mini Kit (Qiagen) and residual DNA was removed with the RNase-Free DNase Kit (Qiagen). A Qubit 2.0 Fluorometer was used to calculate RNA concentration and 4 mg of total RNA was used to prepare libraries with the TruSeq Stranded mRNA Library Prep Kit (Illumina, San Diego, US). Single-end sequencing (50bp) was performed on an Illumina HiSeq 2500. Raw data files have been deposited at NCBI Gene Expression Omnibus (GEO) accession #GSE268259.

### Identification of differentially expressed genes and gene expression analysis

Single end 50bp reads were aligned to the Arabidopsis TAIR10 genome using the HISAT2 pipeline on RMTA (Read Mapping, Transcript Assembly) available through CyVerse’s Discovery Environment [61]. HISAT2 (v2.1.0) was used to align sequencing reads. Reads were assembled into transcripts using StringTie (v1.3.4). Annotation was conducted using TAIR10 FASTA sequence the TAIR10 genome GTF annotation file (www.arabidopsis.org).

The “feature_counts.txt” file obtained from RMTA was used as input for differential gene expression statistical analyses utilizing R studio-DESeq2 available through CyVerse. The DESeq2 was performed between samples (i.e., wt vs. mutants in 24 or cycling light conditions). Genes with a Log_2_ fold-change > +/− 1.5 and a P*adj* < 0.01 were considered differentially expressed genes (DEGs) (**Tables S1-5**).

Heatmaps were generated using ClustalVis on https://biit.cs.ut.ee/clustvis/ [62], and the heat mapper package (https://github.com/WishartLab/heatmapper) on http://www.heatmapper.ca/ [63], using Log_2_ Fold-change values. Average linkage was used for the clustering method and Pearson correlation was used for the distance measurement method. Venn Diagrams comparing the number of DEGs across the four datasets were created using VENNY 2.1 (https://bioinfogp.cnb.csic.es/tools/venny/). To assess the impact of a genotype or condition on certain processes, DEGs were compared to lists of genes involved in senescence (**Table S25**), JA-related processes [64] (**Table S24**) and SA-related processes (**Table S22**), which were manually curated from the literature [32, 65–71]. Hypergeometric tests were used to calculate significance of overlap between gene lists.

Gene Ontology (GO) enrichment analyses were performed using ShinyGO 0.80 (http://bioinformatics.sdstate.edu/go/). The significantly enriched GO terms were determined using default settings and a p-value threshold of ≤ 0.01 (**Tables S6, S7, S10, S11, and S21**).

### Transcription factor analysis

Detailed methods used for transcription factor (TF) analyses may be found in **Supplemental Methods 1**. Briefly, DEGs identified in RNAseq data using a log_2_FC +/− > 1, adjusted p-value ≤ 0.01 cutoff were to identify differentially expressed transcription factors (DETF) and TFs with target genes enriched in these data. Hypergeometric tests were used to identify the enrichment of TF families. Data used to generate heatmaps may be found in **Table S28.** DETFs were identified from DEG lists using PlantTFDB, were compiled (**Table S12 and S13**) and the counts for each family of TF were determined (**Table S14**). GO enrichment analyses were used to identify the enrichment of Biological GO terms represented both in full DETF lists and sub-lists of the different TF families that were significantly enriched in our TF enrichment analysis (**Table S16**). A list of TFs whose targets were overrepresented in DEG lists was obtained from the TF enrichment tool on http://plantregmap.gao-lab.org/tf_enrichment.php (**Table S17**) and the counts for each TF family represented were determined (**Table S14**). Assessment of False Discovery Rate (FDR) for each TF family identified (**Table S18**).

### RNA extraction, cDNA synthesis, and real-time quantitative PCR

Measurement of steady-state RNA transcripts were made as previously described [72] (**Supplemental Methods 2**). **Table S27** lists the primers used.

### DNA extraction and PCR genotyping

To genotype mutant lines, genomic DNA was extracted using a cetyltrimethylammonium bromide-based protocol (Healey et al. 2014). PCR was performed as previously described [15]. **Table S27** lists the primers used.

### Measuring leaf area and chlorophyll content

Leaf area was determined by photographing all leaves in the rosette and calculating total area using ImageJ. Plant chlorophyll content was extracted from total leaf tissue and measured spectrophotometrically according to [73] as previously described [12].

### Cell death measurements

Cell death was measured in adult plants and seedlings using trypan blue staining as previously described [12].

### Leaf senescence assay

The third and fourth rosette leaves from 3-week-old plants were floated on 3 mL of distilled water with or without 100 mM methyl-jasmonate (MeJA). Samples were then incubated in the dark at 21°C for 3 days.

### Jasmonic acid and salicylic acid measurements

SA and JA were extracted from whole rosettes and analyzed as previously described [74, 75]. Plants were grown in 24h constant light for 17 days, and one set of plants were then shifted to 16h light/8h dark diurnal cycling light. Samples were collected two days later, 3hrs after subjective dawn.

### Protein extraction and immunoblot analyses

Total leaf tissue from plants was collected one hour after subjective dawn, frozen in liquid N_2_, and homogenized in extraction buffer (50 mM Tris-Cl pH 8.0, 150 mM NaCl, 0.5% Nonidet P-40 (v:v), and protease inhibitors (Roche)). Cell debris was removed by centrifugation at 10,000 x g for 20 min at 4^0^C. Total protein concentration was measured using the Bradford assay (BioRad) and 15ug of each sample was loaded into a 4-20% SDS-PAGE precast protein gel (BioRad). Proteins were fractionated by electrophoresis and then transferred to PVDF membranes (BioRad). Membranes were incubated with specific primary and secondary antibodies. Immunoblots were imaged by chemiluminescence with the ProSignal Pico ECL Reagents (Genesee Scientific) on a ChemiDoc (Bio-Rad). Polyclonal GFP and Actin antibodies were purchased from BioRad Laboratories. Polyclonal antibodies for NAD(P)H dehydrogenase subunit 45 (NDH45), Thylakoid Membrane Cytochrome B6 Protein (PetB), ATP Synthase Gamma Chain (AtpC), and Rubisco large subunit (RbcL) were purchased from PhytoAB (California, USA). Antibodies for photosystem I core protein (PsaA) and D2 protein of PSII (PsbD) were purchased from Agrisera (Vännäs, Sweden).

### Biomass measurements

Fresh weight was measured in three-week-old plants. For dry weight measurements, samples were stored in envelopes and desiccated at 65°C for 48 hours.

## Supporting information

Suppplemental tables 1-25, 28

## List of abbreviations

JA: Jasmonic acid
PCD: Programmed cell death
ROS: Reactive oxygen species
SA: Salicylic acid
^1^O_2_: Singlet oxygen

## Declarations

## Ethics approval and consent to participate

Not applicable

## Consent for publication

Not applicable

## Availability of data and materials

Raw RNA-seq data and associated metadata files have been deposited at NCBI Gene Expression Omnibus (GEO) (accession #GSE268259) (https://www.ncbi.nlm.nih.gov/geo/query/acc.cgi?acc=GSE268259) and will be made publicly available upon acceptance of this manuscript. All other data is included in the manuscript or Supplementary Material. Further inquiries can be directed to the corresponding author.

## Competing interests

The authors declare that they have no competing interests

## Funding

The authors acknowledge the Division of Chemical Sciences, Geosciences, and Biosciences, Office of Basic Energy Sciences of the U.S. Department of Energy grant DE-SC0019573 awarded to J.D.W., and National Institutes of Health (NIH) grant R01GM107311-8 and National Science Foundation (NSF) grant 2104365 awarded to K.D. M.D.L was supported by the NIH T32 GM136536 training grant and the UA Richard A. Harvill Graduate Fellowship. The funding bodies played no role in the design of the study and collection, analysis, and interpretation of data and in writing the manuscript.

## Authors’ contributions

SR and JDW planned and designed the research. SR performed RNA-Seq data analysis, created higher-order mutants, performed genotyping, phenotyping, biochemical assays, cell death measurements, and immunoblotting. MDL assisted with processing RNA-seq data, performed transcription factor analyses. AMA assisted with genotyping and the leaf senescence assay. MFGM and KD measured JA and SA content. JDW conceived the original scope of the project and managed the project. SR, MDL, MFGM, KD, and JDW contributed to data analysis and interpretation, and reviewed the manuscript. SR, MDL, and JDW wrote the manuscript. All authors approved the final version.

## Acknowledgments

The authors thank Kamran Alamdari (University of Arizona) for technical assistance with RNA extractions and RT-qPCR analyses, and Sophia Daluisio and Alexa Abate (University of Arizona) for technical assistance with plant maintenance.

## Notes

### Competing Interest Statement

The authors have declared no competing interest.

### Summary of Updates

This revised version (v2) has been altered to focus on the key findings and analyses of the included RNA-seq dataset. This included major revisions to the introduction and discussion sections to focus on key findings and the necessary background. Additional analyses on the expression of microautophagy and senescence pathway genes were included. Additional statistical analyses for gene enrichment studies were included. Finally, experiments and figure panels involving the genetic analyses of hormone synthesis mutants has been removed to focus on the core results and conclusions from the RNA-seq dataset analyses. Overall, this has led to a more streamlined presentation of the data and conclusions.

