## Supplemental materials (figures, methods, tables) for "Transcript profiling of *plastid ferrochelatase two* mutants reveals that chloroplast singlet oxygen signals lead to global changes in RNA profiles and are mediated by Plant U-Box 4"

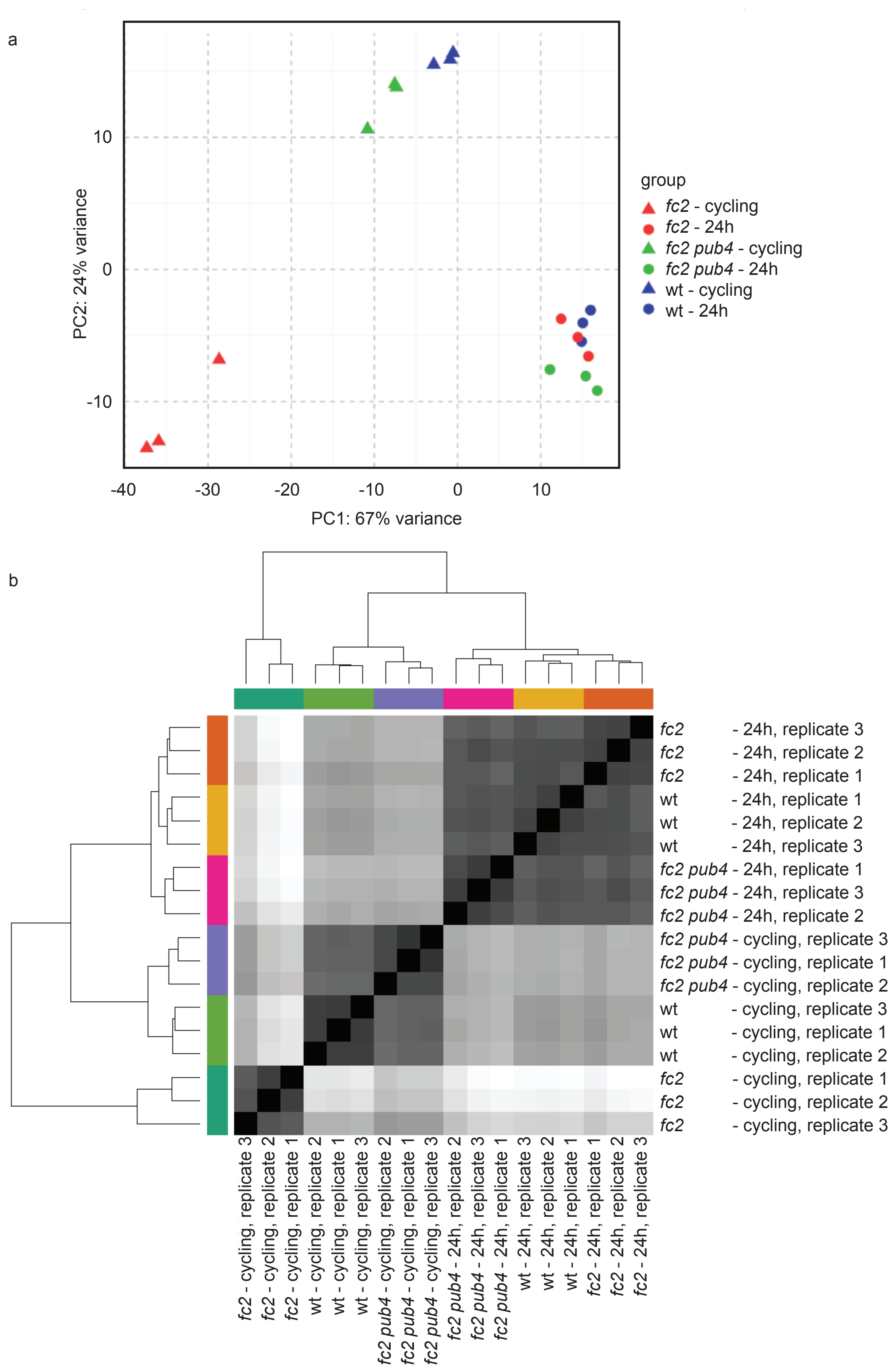


Figure S1. Visualization of variance among RNA-seq replicates used in the study.

Shown are two analyses to visualize variance among the 18 RNA-seq replicates used in this study. Three genotypes (wt, *fc2*, and *fc2 pub4*) were grown in two conditions (19 days in constant light conditions (24h) or 17 days of constant light conditions followed by two days on 16h light/8h dark diurnal cycling light conditions (cycling)) for a total of 6 groups with three replicates each. (a) Principle component analysis (PCA) scatter plot showing variance of the RNA-seq replicates. Principle component (PC) 1 and PC2 are plotted and colored by genotype (wt, *fc2* and *fc2 pub4*) and growing condition (constant light (24h) and 16h light/8h dark diurnal cycling light (cycling) conditions) for each replicate (3 each). The genotypes and conditions are color coded according to the key on the right. PCA was performed using feature count matrix produced from the RNA-seq data sets. Variation percentage for each PC (PC1: 67% variance, PC2: 24% variance) reported in brackets with the axis label. (b) Shown is a distance matrix for the same 18 replicates. The heatmap was generated using feature count matrix produced from the RNA-seq data sets and the clustering of samples show an overview of similarities and dissimilarities between the replicates. Shades of grey indicate distance between samples with darker shades indicating smaller distance and lighter shades indicating greater distance.

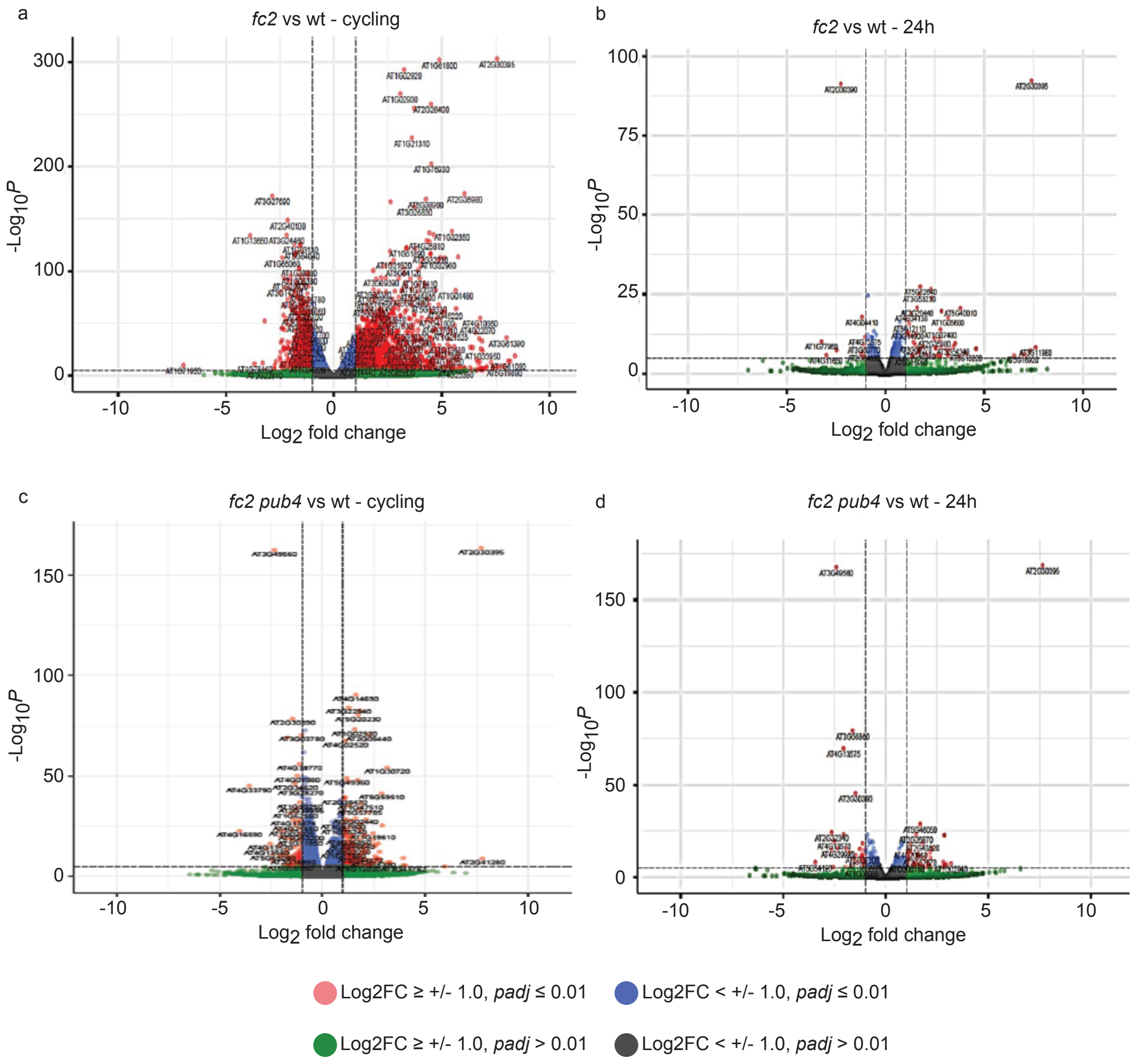


<u>Figure S2. Volcano plot analysis of mean differential gene expression between genotypes in a single condition.</u>

Volcano plots showing pairwise comparisons between genotypes (wt vs. *fc2* or wt vs. *fc2 pub4*) within a single condition (permissive constant light (24h) conditions or two days of singlet oxygen-producing 16h light/8h dark diurnal cycling light conditions (cycling)). Expression data is from the DESeq2 analysis of the included RNA-seq data set; (a) *fc2* vs. wt in cycling light conditions, (b) *fc2* vs. wt in 24h light conditions, (c) *fc2 pub4* vs. wt in cycling light conditions, and (d) *fc2 pub4* vs. wt in 24h light conditions. The y-axis represents the negative $\log_{10}$-transformed *P*-values from gene-specific tests, and the x-axis shows the $\log_2$ fold change (FC). Red dots represent differentially expressed genes (DEGs) according to the $\log_2$FC and *padj* cut-off values of ≥ +/-1 and ≤ 0.01 respectively. Green dots represent DEGs that only pass the $\log_2$FC cutoff, blue dots represent DEGs that only pass the $\log_2$FC cutoff, and dark grey dots represent DEGs that did not pass either cutoff. The most upregulated genes are towards the right, the most downregulated genes are towards the left, and the most statistically significant genes are towards the top.

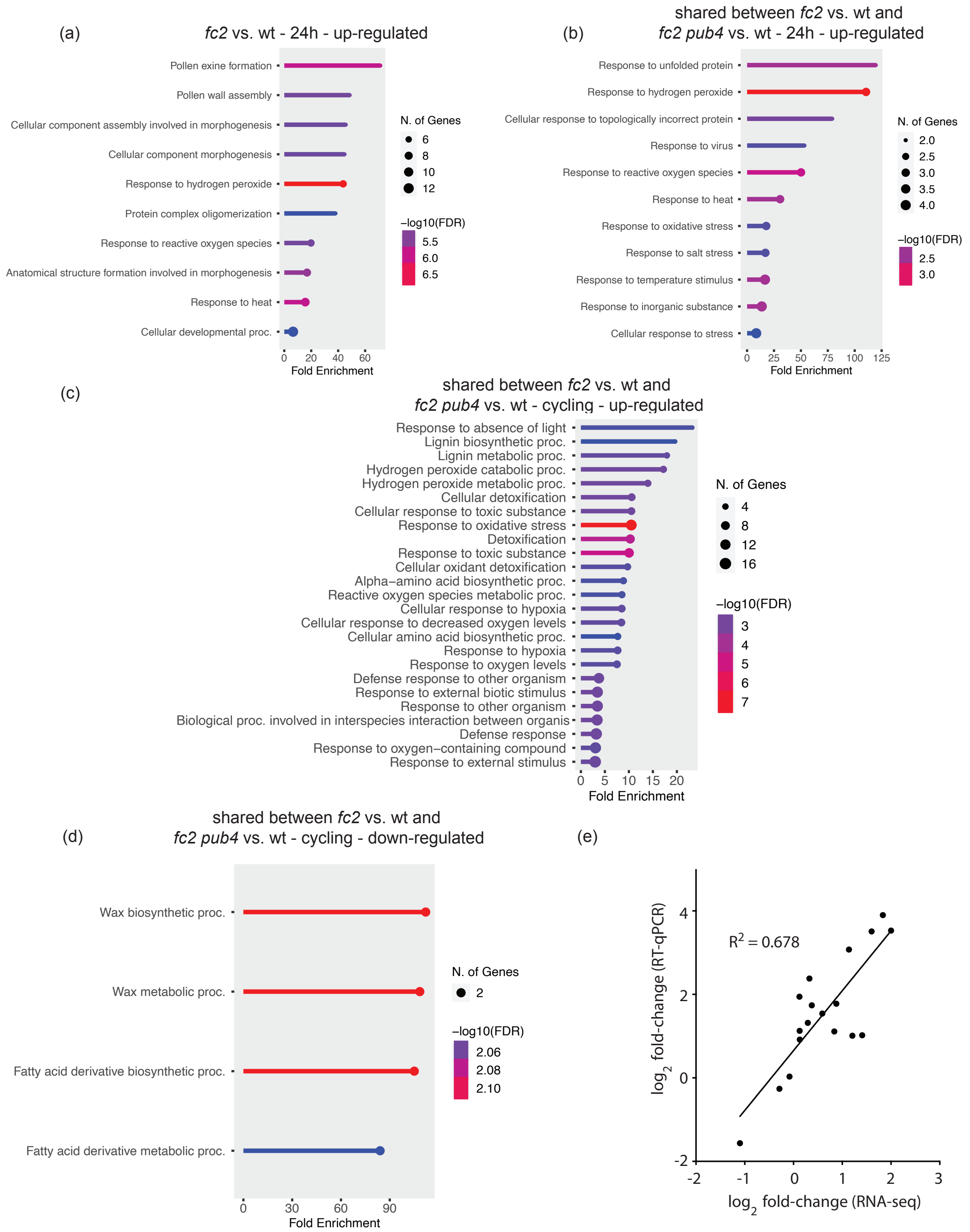


<u>Figure S3. Gene ontology analyses of differentially expressed genes.</u>

Shown are gene ontology (GO) term enrichment analyses of the identified differentially expressed genes (DEGs) in this study. (a) GO-term enrichment from the 60 up-regulated DEGs from *fc2* vs.

wt under constant (24h) light conditions. (b) GO-term enrichment from the 11 shared up-regulated DEGs between *fc2* vs. wt and *fc2 pub4* under constant (24h) light conditions. (c) GO-term enrichment from the 80 shared up-regulated DEGs between *fc2* vs. wt and *fc2 pub4* under 16h light/8h dark diurnal cycling light (cycling) conditions. (d) GO-term enrichment from the 80 shared down-regulated DEGs between *fc2* vs. wt and *fc2 pub4* under 16h light/8h dark diurnal cycling light (cycling) conditions. GO-term enrichment analyses were performed using ShinyGO 0.80 with an FDR cutoff off of $p \leq 0.01$. x-axes indicate fold-enrichment. The ball size indicates number of genes. The line colors represent FDR values. (e) RNA-seq analysis of plants grown in cycling light conditions was confirmed by RT-qPCR using RNA extracted from an independent set of plants grown in the same conditions (17 days of 24h constant light conditions, followed by two days of 16h light/8h dark diurnal cycling light conditions and harvested one hour post dawn). Shown is the correlation of independent RNA-seq and RT-qPCR analyses of pairwise comparisons between wt and *fc2* under cycling light conditions and wt and *fc2 pub4* under cycling light conditions. The fold change values (log2) of nine nuclear genes (*SIB1*, *PAO*, *CHLH*, *LOX4*, *AOC3*, *EDS16*, *CAM1*, *GPA1*, *PAP1*) are plotted (total of 18 comparisons). Expression values normalized to expression of *ACTIN2*.

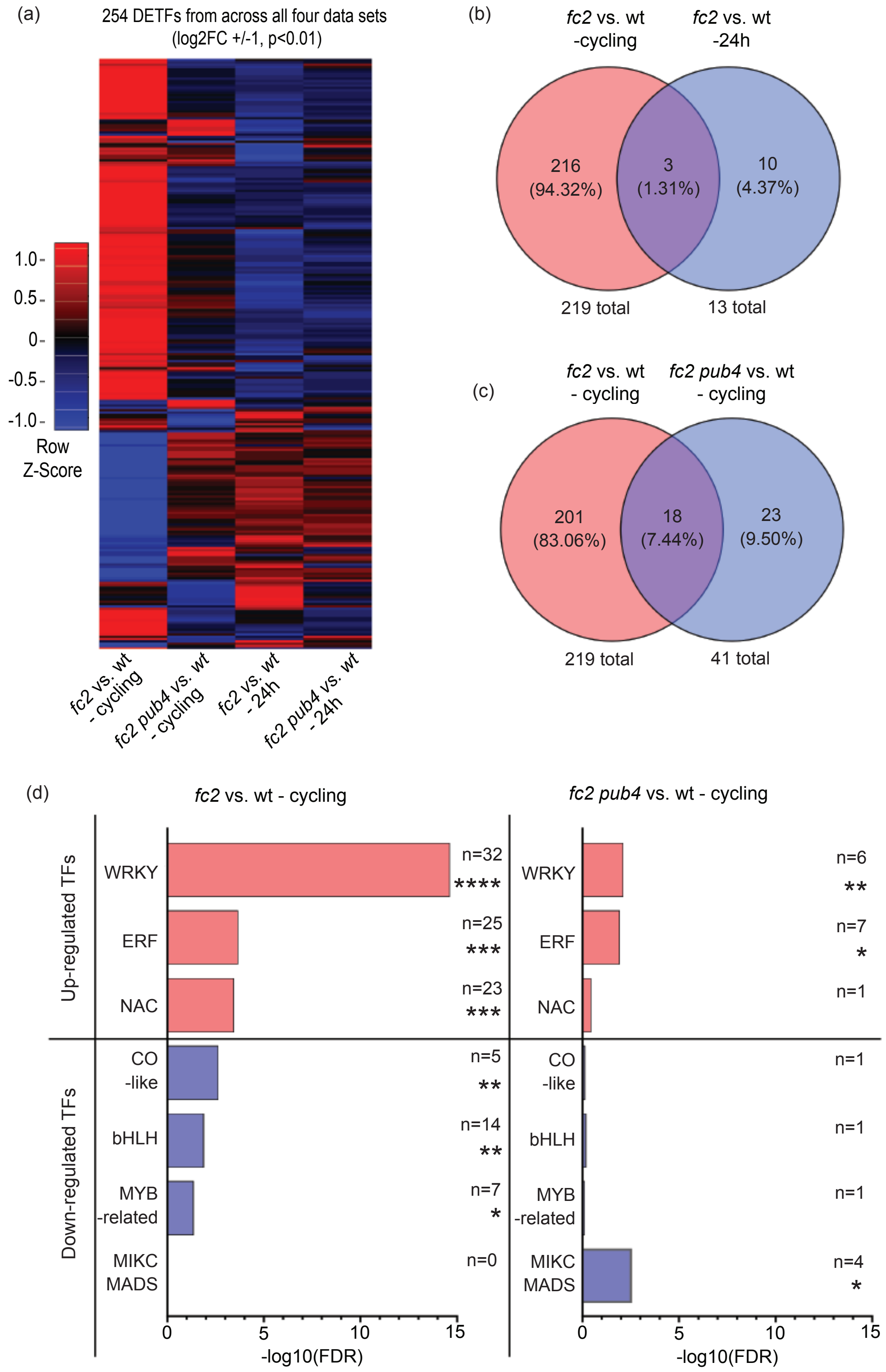


Figure S4. WRKY, ERF, and NAC transcription factor networks are induced by chloroplast singlet oxygen accumulation.

The RNA-seq data set was analyzed for differentially expressed transcription factors (DETFs) (a) A heatmap showing the change in expression of 254 DETFs identified from all differentially expressed genes (DEGs) lists from pairwise comparisons between wt and the mutants (*fc2* or *fc2 pub4*) in each condition (constant light conditions (24h) or 16h light/8h dark diurnal cycling light conditions (cycling) (cutoffs ≥ +/- 1.0 $\log_2$FC, *padj* ≤ 0.01). Red color indicates up-regulation, blue indicates down-regulation, and black indicates no change. (b) Venn diagram comparing the DETFs identified in DEG lists obtained from "*fc2* vs. wt – cycling" (219 genes) and "*fc2* vs. wt - 24h" (13 genes). (c) Venn diagram comparing the DETFs identified in DEG lists obtained from "*fc2* vs. wt – cycling" (219 genes) and "*fc2 pub4* vs wt – cycling" (41 genes). (d) Enrichment of up-regulated (red bars) and down-regulated (blue bars) DETF families from "*fc2* vs. wt – cycling" and "*fc2 pub4* vs. wt – cycling" analyses. n represents the count of respective TF family genes represented in each list. A hypergeometric test followed by the Benjamini–Hochberg method was used to determine the False Discovery Rate (FDR) (* = $P \leq 0.05$, ** = $P \leq 0.01$, *** = $P \leq 0.001$, **** = $P \leq 0.0001$).

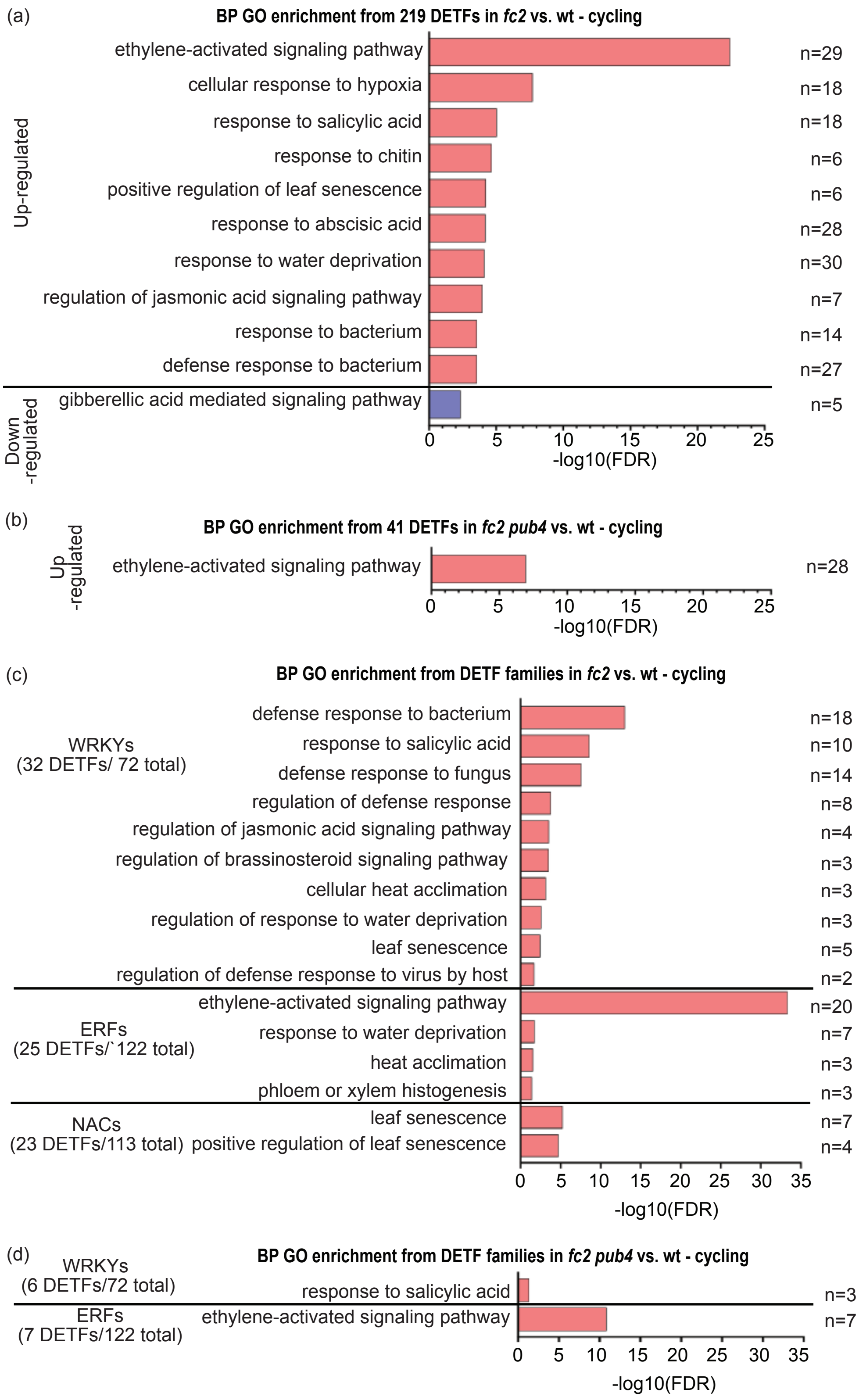


Figure S5. Gene Ontology term enrichment of differentially expressed transcription factors genes and transcription factor gene families.

Differentially expressed transcription factor (DETF) and DETF families from the "*fc2* vs. wt – cycling" (219 genes) and "*fc2 pub4* vs. wt – cycling" (41 genes) pairwise comparisons were assessed for biological process (BP) gene ontology (GO) term enrichment. (a) BP GO term enrichment from the "*fc2* vs. wt – cycling" pairwise comparison (219 genes). (b) BP GO terms enrichment from the "*pub4* vs. wt -cycling" pairwise comparison (41 genes). (c) BP GO term enrichment of WRKY (32 genes), ERF (25 genes), and NAC (23 genes) transcription factor (TF) family genes up-regulated in the "*fc2* vs. wt – cycling" pairwise comparison (d) BP GO term enrichment of WRKY (32 genes) and ERF (25 genes) TF family genes up-regulated in the "*fc2 pub4* vs. wt – cycling" pairwise comparison. Red bars = up-regulated and blue bars = down-regulated, respectively. GO term enrichment analysis was performed using the Database for Annotation, Visualization, and Integrated Discovery (DAVID) 2021 (https://david.ncifcrf.gov/). Top 10 significant terms (where applicable) are reported. n represents the count of respective TF or TF family genes represented in each list. An FDR ≤ 0.05 cutoff was applied to enriched GO terms found in this analysis.

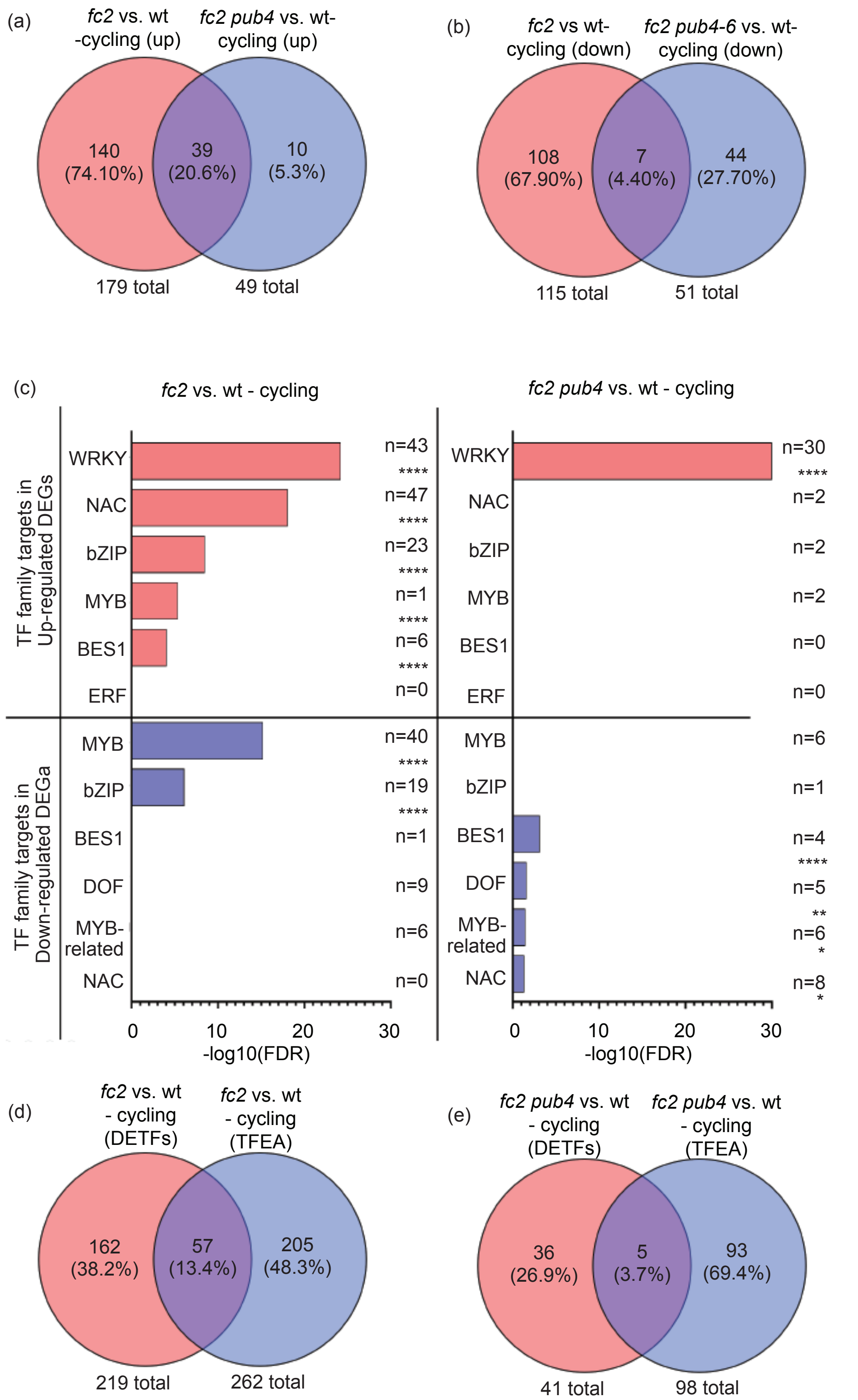


Figure S6. WRKY and NAC transcription factor targets are induced by chloroplast singlet oxygen accumulation and activation of NAC targets is blocked by the *pub4* mutation.

Transcript levels determined by RNA-seq analyses was used to determine the transcription factors (TFs) whose targets were enriched in chloroplast singlet oxygen ($^1O_2$) signaling. Venn diagrams depicting the transcription factor enrichment analyses (TFEAs) which identified the TFs with enriched targets in (a) up-regulated and (b) down-regulated genes identified in differentially expressed gene (DEG) lists obtained from "*fc2* vs. wt – cycling" and "*fc2 pub4* vs. wt – cycling" pairwise comparisons. (c) Enrichment of TF families from "*fc2* vs. wt – cycling" and "*fc2 pub4* vs. wt – cycling" pairwise comparisons whose targets were significantly enriched in their corresponding DEG lists. Red bars = up-regulated and blue bars = down-regulated, respectively. n represents the count of respective TF family genes represented in each list. A hypergeometric test followed by the Benjamini–Hochberg method was used to determine the False Discovery Rate (FDR) (* = $P \leq 0.05$, ** = $P \leq 0.01$, *** = $P \leq 0.001$, **** = $P \leq 0.0001$). Venn diagram comparing the DETFs identified in **Figs. S4b and c** with TFs obtained from the TFEA using up- and down-regulated DEGs from (d) "*fc2* vs. wt – cycling" (262 TFs) and (e) "*fc2 pub4* vs. wt – cycling" (98 TFs) pairwise comparisons.

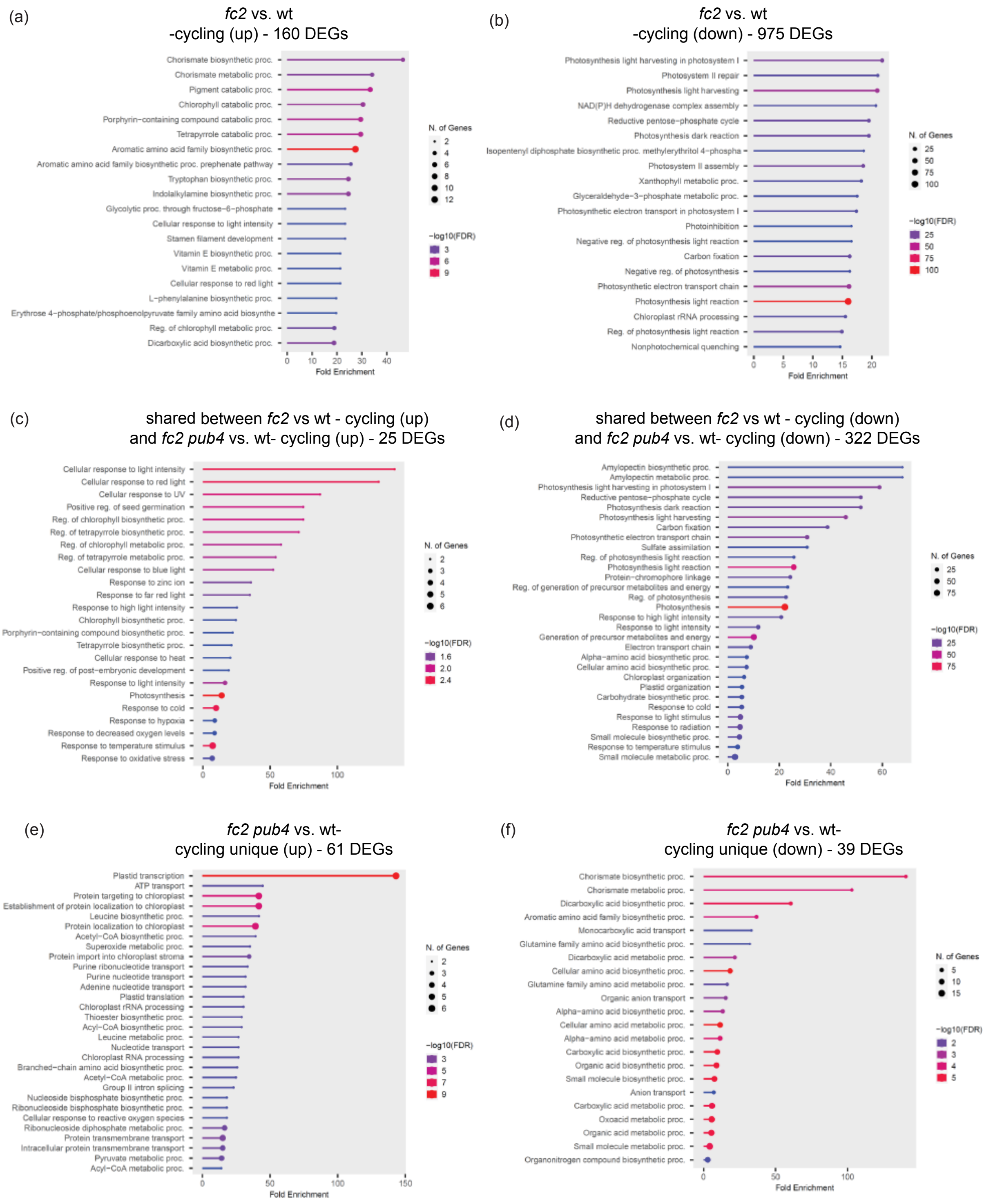


<u>Figure S7. Gene ontology analyses of differentially expressed plastid protein-encoding genes.</u> Shown are gene ontology (GO) term enrichment analyses of the identified differentially expressed plastid protein encoding genes (PPEGs) in this study. (a) GO-term enrichment from the 160 up-regulated PPEGs from *fc2* vs. wt under 16h light/8h dark diurnal cycling light conditions (cycling).

(b) GO-term enrichment from the 975 down-regulated PPEGs from *fc2* vs. wt - cycling. (c) GO-term enrichment from the shared 25 up-regulated PPEGs between *fc2* vs. wt – cycling and *fc2 pub4* – cycling. (d) GO-term enrichment from the shared 322 down-regulated PPEGs between *fc2* vs. wt – cycling and *fc2 pub4* – cycling. (e) GO-term enrichment from the unique 61 up-regulated PPEGs from *fc2 pub4* – cycling. (f) GO-term enrichment from the unique 39 down-regulated PPEGs from *fc2 pub4* – cycling. GO-term enrichment analyses were performed using ShinyGO 0.80 with an FDR cutoff off of $p \leq 0.01$. x-axes indicate fold-enrichment. The ball size indicates number of genes. The line colors represent FDR values.

(a)

| Plastid encoded chloroplast gene expression (Log2FC values) | | | | |
|---|---|---|---|---|
| | | | *fc2* vs. wt - cycling | *fc2 pub4* vs. wt - cycling |
| | *ATCG00350* | *psaA* | -0.16 | 0.12 |
| PSI | *ATCG00340* | *psaB* | -0.42 | -0.17 |
| | *ATCG01060* | *psaC* | 0.22 | 0.21 |
| | *ATCG00510* | *psaI* | -0.52 | -0.47 |
| | *ATCG00630* | *psaJ* | -0.86 | -0.16 |
| | *ATCG00020* | *psbA* | 0.09 | -0.32 |
| | *ATCG00680* | *psbB* | 0.19 | -0.09 |
| | *ATCG00280* | *psbC* | 0.33 | 0.12 |
| | *ATCG00270* | *psbD* | 0.24 | -0.10 |
| | *ATCG00580* | *psbE* | -0.29 | 0.04 |
| PSII | *ATCG00570* | *psbF* | 1.94 | 1.36 |
| | *ATCG00430* | *psbG* | -0.13 | 0.02 |
| | *ATCG00710* | *psbH* | 0.37 | 0.36 |
| | *ATCG00080* | *psbI* | -1.23 | -0.96 |
| | *ATCG00550* | *psbJ* | -0.10 | -0.23 |
| | *ATCG00070* | *psbK* | -1.03 | -0.49 |
| | *ATCG00560* | *psbL* | -0.25 | 0.52 |
| | *ATCG00220* | *psbM* | 0.08 | 0.76 |
| | *ATCG00700* | *psbN* | 0.37 | 0.24 |
| | *ATCG00690* | *psbT* | -0.55 | 0.44 |
| | *ATCG01100* | *ndhA* | 0.21 | 0.07 |
| | *ATCG00440* | *ndhC* | -1.22 | -1.17 |
| | *ATCG01050* | *ndhD* | -0.74 | -0.14 |
| NDH | *ATCG01070* | *ndhE* | -0.19 | -0.20 |
| complex | *ATCG01010* | *ndhF* | -0.73 | -0.21 |
| | *ATCG01080* | *ndhG* | -0.68 | -0.30 |
| | *ATCG01110* | *ndhH* | 0.13 | 0.58 |
| | *ATCG01090* | *ndhI* | 0.07 | 0.29 |
| | *ATCG00420* | *ndhJ* | -0.68 | -0.39 |
| | *ATCG00120* | *atpA* | -0.76 | -0.33 |
| | *ATCG00480* | *atpB* | -0.85 | -0.59 |
| ATP | *ATCG00470* | *atpE* | -0.75 | -0.38 |
| synthase | *ATCG00130* | *atpF* | -0.24 | -0.13 |
| | *ATCG00140* | *atpH* | 0.27 | -0.33 |
| | *ATCG00150* | *atpI* | 0.05 | 0.23 |
| | *ATCG00540* | *petA* | -0.72 | -0.21 |
| Cyt b6f | *ATCG00720* | *petB* | -0.23 | 0.00 |
| complex | *ATCG00730* | *petD* | -0.18 | -0.19 |
| | *ATCG00600* | *petG* | -1.49 | 0.09 |
| RUBISCO | *ATCG00490* | *rbcL* | -0.35 | -0.25 |

(b)

| Nuclear encoded chloroplast gene expression (Log2FC values) | | | | |
|---|---|---|---|---|
| | | | *fc2* vs. wt - cycling | *fc2 pub4* vs. wt - cycling |
| | *AT4G02770* | *psaD1* | -1.32 | -0.29 |
| | *AT1G03130* | *psaD2* | -1.55 | -0.46 |
| | *AT4G28750* | *psaE1* | -1.43 | -0.37 |
| | *AT2G20260* | *psaE2* | -1.19 | -0.10 |
| PSI | *AT1G31330* | *psaF* | -1.42 | -0.37 |
| | *AT1G55670* | *psaG* | -1.43 | -0.40 |
| | *AT3G16140* | *psaH1* | -1.22 | -0.23 |
| | *AT1G52230* | *psaH2* | -1.59 | -0.48 |
| | *AT1G30380* | *psaK* | -1.61 | -0.37 |
| | *AT4G12800* | *psaL* | -1.43 | -0.35 |
| | *AT5G64040* | *psaN* | -1.49 | -0.44 |
| | *AT1G08380* | *psaO* | -1.59 | -0.46 |
| | *AT2G46820* | *psaP* | -1.42 | -0.52 |
| | *AT5G66570* | *psbO1* | -1.11 | -0.27 |
| | *AT3G50820* | *psbO2* | -0.97 | -0.23 |
| | *AT1G06680* | *psbP1* | -1.29 | -0.38 |
| | *AT2G30790* | *psbP2* | -0.80 | -0.61 |
| | *AT4G21280* | *psbQ1* | -1.55 | -0.53 |
| PSII | *AT4G05180* | *psbQ2* | -1.60 | -0.42 |
| | *AT1G79040* | *psbR* | -1.06 | -0.12 |
| | *AT1G44575* | *psbS* | -1.08 | -0.24 |
| | *AT3G21055* | *psbTN* | -1.40 | -0.16 |
| | *AT2G30570* | *psbW* | -1.16 | -0.31 |
| | *AT2G06520* | *psbX* | -1.19 | -0.35 |
| | *AT1G67740* | *psbY* | -1.13 | -0.28 |
| | *AT1G15980* | *ndh48* | -1.14 | -0.38 |
| NDH | *AT1G64770* | *ndh45* | -0.94 | -0.13 |
| complex | *AT5G43750* | *ndh18* | -1.38 | -0.41 |
| | *AT3G16250* | *ndf4* | -1.12 | -0.36 |
| | *AT1G18730* | *ndf6* | -1.28 | -0.34 |
| ATP | *AT4G04640* | *atpC1* | -0.96 | -0.26 |
| synthase | *AT4G09650* | *atpD* | -1.16 | -0.46 |
| | *AT4G03280* | *petC* | -1.14 | -0.20 |
| Cyt b6f | *AT1G76100* | *petE1* | -1.15 | -0.26 |
| complex | *AT1G20340* | *petE2* | -1.30 | -0.31 |
| | *AT2G26500* | *petM* | -0.93 | -0.14 |
| | *AT3G54890* | *lhca1* | -1.66 | -0.38 |
| | *AT3G61470* | *lhca2* | -1.47 | -0.32 |
| | *AT1G61520* | *lhca3* | -1.48 | -0.42 |
| | *AT3G47470* | *lhca4* | -1.71 | -0.53 |
| | *AT1G19150* | *lhca6* | -1.58 | -0.46 |
| | *AT1G29920* | *lhcb1.2* | -2.03 | -0.28 |
| Light | *AT2G34430* | *lhcb1.4* | -3.29 | -0.78 |
| harvesting | *AT2G05070* | *lhcb2.2* | -1.84 | -0.33 |
| complex | *AT3G27690* | *lhcb2.3* | -2.85 | -0.71 |
| | *AT5G54270* | *lhcb3* | -1.81 | -0.58 |
| | *AT5G01530* | *lhcb4.1* | -1.45 | -0.37 |
| | *AT3G08940* | *lhcb4.2* | -1.83 | -0.58 |
| | *AT2G40100* | *lhcb4.3* | -2.13 | -0.10 |
| | *AT4G10340* | *lhcb5* | -1.71 | -0.53 |
| | *AT1G15820* | *lhcb6* | -1.83 | -0.59 |

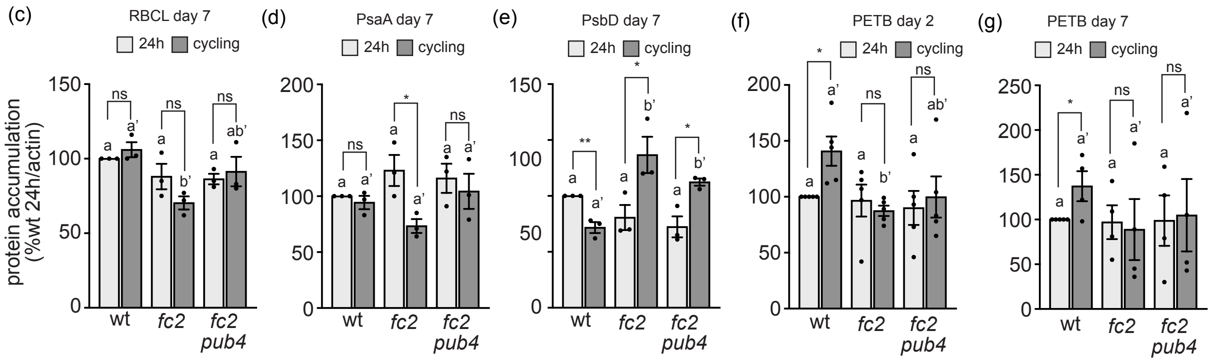


Figure S8. Relative expression of plastid protein-encoding genes during singlet oxygen stress.

Relative expression of select (a) plastid-encoded and (b) nuclear encoded (right column) chloroplast proteins in *fc2* and *fc2 pub4,* grown in 16h light/8h dark diurnal (cycling) light conditions, compared to wt. Shown are $Log_2$ Fold Change values from the DESeq2 analysis of the included RNA-seq data set. Values shaded in blue pass the significance cutoff (padj ≤ 0.01), and values in red font do not. The left columns indicate the protein complexes in which these gene products reside. Immunoblot analysis of selected plastid-encoded proteins (c) RbcL, (d) PsaA, (e) PsbD, and PetB (f and g) from leaves of three-week-old plants grown in constant (24h) light conditions or grown in 24h light conditions and shifted to 16h light/8h dark diurnal (cycling) light conditions for two (f) or seven (c, d, e, and g) days. Shown are mean values compared to actin levels (+/- SEM) and normalized to wt in 24h light conditions (n ≥ 3 leaves from separate plants). Statistical analyses were performed using one-way ANOVA tests, and the different letters above the bars indicate significant differences within data sets determined by Tukey-Kramer post-tests (P ≤ 0.05). Separate analyses were performed for the different light conditions, and the significance for the cycling light condition is denoted by letters with a prime symbol (′). Statistical analyses of genotypes between conditions were performed by student's t-tests (*, $P \leq 0.05$; ns, P ≥ 0.05). Closed circles indicate individual data points.

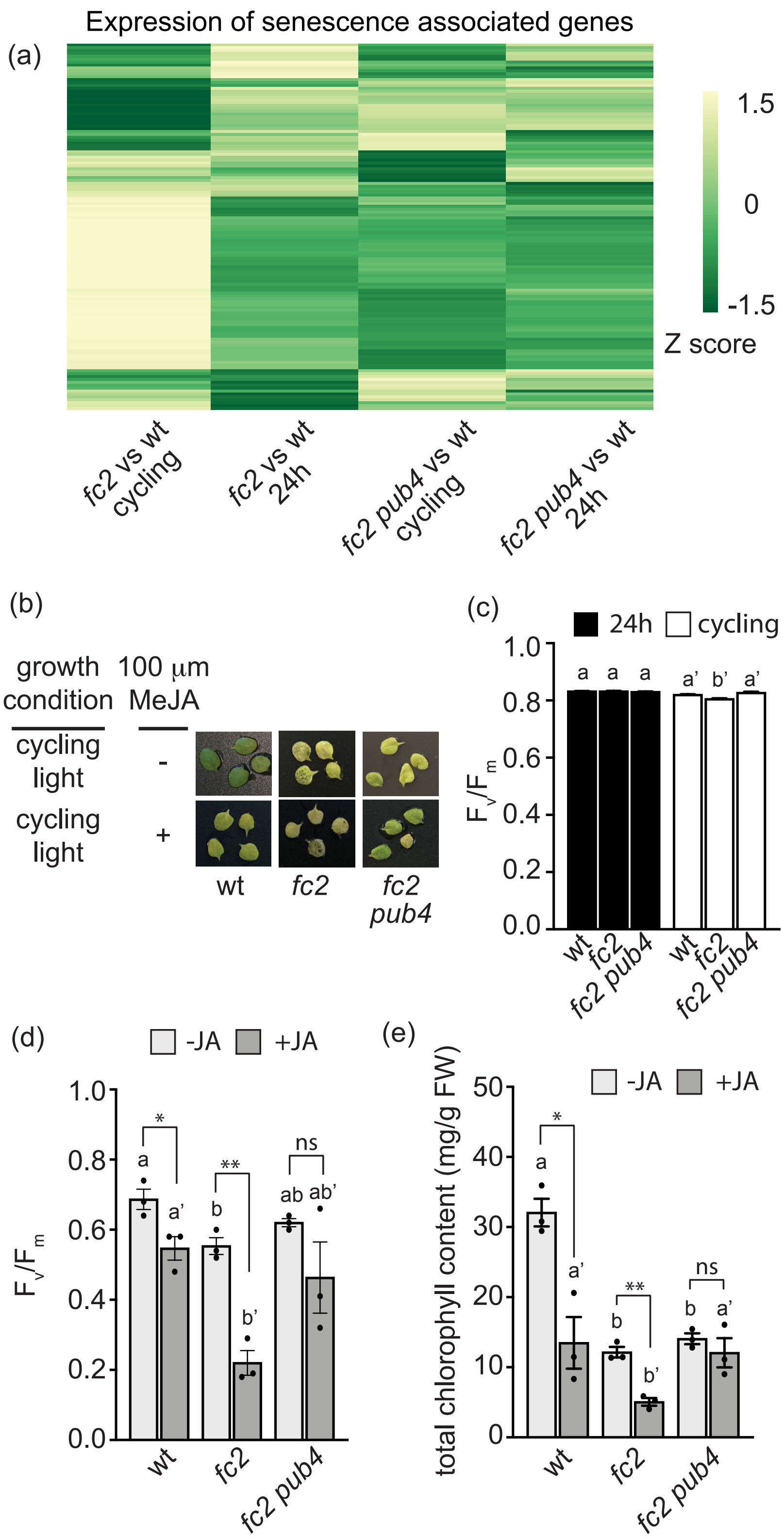


<u>Figure S9. Activation of senescence pathways is blocked by the *pub4* mutation.</u>

(a) Heatmap showing the expression levels of 128 senescence-associated genes (GO-term "leaf senescence," GO:0010150) in *fc2* and *fc2 pub4* (relative to the wt), under constant (24h) light and 16h light/8h dark (cycling) light conditions. The gene expression data are $Log_2$ Fold Change values derived from DESeq2 analysis of the included RNA-seq data set. Genes that are up-regulated or down-regulated in comparison to wt are colored yellow or green, respectively. (b) Representative images of detached 3rd and 4th rosette leaves from plants grown in 24h light conditions plus two days of cycling light conditions. The leaves were incubated in the dark with water (control conditions) or with 100 μM methyl jasmonate (MeJA) for 3 days. Measurement of the maximum quantum efficiency of photosystem II ($F_v/F_m$) of (c) control leaves (left in light without JA) or (d) leaves from panel b. (e) Total chlorophyll content of leaves in panel b. Statistical analyses were performed using one-way ANOVA tests, and the different letters above the bars indicate significant differences within data sets determined by Tukey-Kramer post-tests ($P \leq 0.05$). Separate analyses were performed in c for each light condition and in d and e for the different treatments. The significance for cycling light samples in c and for the JA treatment in d and e are denoted by letters with a prime symbol (′). Statistical analyses of genotypes between conditions were performed by student's t-tests (*, $P \leq 0.05$; **, $P \leq 0.01$; ns, $P \geq 0.05$). n = 3 whole leaves from separate plants. Error bars = +/- SEM. Closed circles indicate individual data points.

**Methods S1.** Transcription factor analysis

*Data used in TF analysis.*

To perform transcription factor (TF) enrichment analyses, DESeq2 was used to identify differentially expressed genes (DEGs) (from all four pairwise comparisons: *fc2* vs. wt – 24h, *fc2* vs. wt – cycling, *fc2 pub4* vs. wt – 24h, *fc2 pub4* vs. wt – cycling) with the following cutoff criteria: $log_2FC$ +/- ≥ 1, adjusted p-value ≤ 0.01. DEGs that passed these cutoff criteria (3121 genes) were compared to a list of known transcription factors in Arabidopsis (representing 1717 unique loci) from PlantTFDB to identify 254 TFs (http://planttfdb.gao-lab.org/index.php?sp=Ath) [2-4].

*Heatmap generation.*

The expression of these 254 genes were compared across all four pairwise data sets to produce an expression matrix (**Table S28**, which was used to generate a heatmap using the heat mapper package (https://github.com/WishartLab/heatmapper) on http://www.heatmapper.ca/ [5].

*TF enrichment.*

Differentially expressed TFs (DETFs) were identified from DEG lists using PlantTFDB, were compiled (**Table S11 and S12**) and the counts for each family of TF were determined (**Table S13**). A hypergeometric test followed by the Benjamini–Hochberg method was used to determine the False Discovery Rate (FDR) of each TF family identified (**Table S14**). A cutoff (FDR < 0.05) was applied and the -log10(FDR) was calculated for each TF family that passed this cutoff.

*TF Gene Ontology Analysis*. Gene ontology analysis was performed using Database for Annotation, Visualization and Integrated Discovery (DAVID) 2021 (https://david.ncifcrf.gov/) [6, 7]. Lists of DETFs from cycling light-exposed plants (*fc2* vs. wt - cycling and *fc2 pub4* vs. wt - cycling) were used to identify the enrichment of Biological GO terms represented both in full DETF lists and sub lists of the different TF families that were significantly enriched in our TF enrichment analysis (**Table S15**). The reported GO terms represent the top 10 statistically significant (FDR ≤ 0.05) GO terms excluding terms relating to transcriptional regulation.

*Transcription factor enrichment analysis (TFEA).*

A list of TFs whose targets were overrepresented in DEG lists was obtained from the TF enrichment tool on http://plantregmap.gao-lab.org/tf_enrichment.php (**Table S16**) and the counts for each TF family represented were determined (**Table S13**). A hypergeometric test followed by the Benjamini–Hochberg method was used to determine the False Discovery Rate (FDR) of each TF family identified (**Table S17**). A cutoff (FDR ≤ 0.05) was applied and the -log10(FDR) was calculated for each TF family that passed this cutoff.

**Methods S2.** RNA extraction, cDNA synthesis, and real-time quantitative PCR

Measurement of steady-state RNA transcripts were made as previously described [8]. The RNeasy Plant Mini Kit (Qiagen) was used to extract total RNA from whole seedlings. Next, cDNA was synthesized using the Maxima first strand cDNA synthesis kit for RT-qPCR with DNase (Thermo Scientific) following the manufacturer's instructions. Real-time PCR was performed using the SYBR Green Master Mix (BioRad) with the SYBR Green fluorophore and a CFX Connect Real Time PCR Detection System (BioRad). The following 2-step thermal profile was used in all RT-qPCR: 95 °C for 3 min, 40 cycles of 95 °C for 10s and 60 °C for 30s. *ACTIN2* expression was used as a standard to normalize all gene expression data. **Table S27** lists the primers used.

Table S26. Mutant lines used in this study

| Mutant | Gene | mutation | notes | ref |
|---|---|---|---|---|
| *fc2-1* | *PLASTID FERROCHELATASE 2, FC2, AT2G30390* | GABI_766H08<br>T-DNA in 5'UTR | Sulfadiazin$^r$ | [9] |
| *aos1-1 /dde2* | *ALLENE OXIDE SYNTHASE, AOS, DELAYED DEHISCENCE 2, DDE2, AT5G42650* | SALK_017756<br>T-DNA in exon | | [10] |
| *eds16-1 /sid2-2* | *ISOCHORISMATE SYNTHASE 1, ENHANCED DISEASE SUSCEPTIBILITY TO ERYSIPHE ORONTII 16, EDS16, SALICYLIC ACID INDUCTION DEFICIENT 2, SID2, AT1G74710* | Deletion/rearrangement in exon 9. | | [11] |
| *pub4-6* | *PLANT U-BOX 4, PUB4, AT2G23140* | Point mutation (c9847535t) leading to amino acid substitution (G255R) | | [12] |

Table S27. Primers used in study

| | | |
|---|---|---|
| **RT-qPCR primers** | | |
| *AT3G18780 (ACTIN2)* | For. JP199 | GCACTTGCACCAAGCAGCAT |
| | Rev. JP200 | CCTTTCAGGTGGTGCAACGAC |
| *AT3G56710 (SIB1)* | For. JP589 | CAACCGGAGCCCATCTATT |
| | Rev. JP590 | GGAGAAAGGTTGTGGTCGTC |
| *AT3G44880 (PAO)* | For. WLO1690 | TTGGATCCGAATGTGCCAAC |
| | Rev. WLO1691 | TCAAAGGCTGCCCATTTCTG |
| *AT5G13630 (CHLH)* | For. WLO1692 | TGCAAGCTTTGGAAGGCAAG |
| | Rev. WLO1693 | ACTTGCCATTGCTGCTGTTG |
| *AT1G72520 (LOX4)* | For. WLO1696 | TTGCGTCTCCGTTTCATTCG |
| | Rev. WLO1697 | ACAACGCCGCTTGAGTTAAC |
| *AT3G25780 (AOC3)* | For. WLO1698 | ACTCGGCAAGAAACCAACAG |
| | Rev. WLO1699 | TGATTCCCACGCGTTTCTTG |
| *AT1G74710 (EDS16)* | For. WLO1704 | TGGTGCACCAGCTTTTATCG |
| | Rev. WLO1705 | AAAGCTTCACTGCAGACACC |
| *AT5G37780 (CAM1)* | For. WLO1712 | ATGGAAACGGCACTATCGAC |
| | Rev. WLO1713 | TTGTCGAAAACCCTGAAGGC |
| *AT2G26300 (GPA1)* | For. WLO1714 | TGCAAGAGTTCGCACAACTG |
| | Rev. WLO1715 | TCCACCCACGTCAAACAATC |
| AT1G56650 (*PAP1*/*MYB75*) | For. WLO1959 | GCGAAAAGGTGCTTGGACTAC |
| | For. WLO1960 | ACTTGGTGCCATTTGCCTTC |
| *AT2G14610* (*PR1*) | For. WLO1771 | GTGCTCTTGTTCTTCCCTCGA |
| | Rev. WLO1772 | CCCACGAGGATCATAGTTGCA |
| *AT2G38470* (*WRKY33*) | For. WLO1632 | CCAAACCGAGACTCGTCCAA |
| | Rev. WLO1633 | TGCACTACGATTCTCGGCTC |
| *AT1G80840* (*WRKY40*) | For. WLO1812 | ACAACCATCCAATGCCATCG |
| | Rev. WLO1813 | TCTTCTGTTTGCTGCAACGG |
| *AT3G52430* (*PAD4*) | For. WLO1694 | ACGAGTGAGTTGCAAGCTTC |
| | Rev. WLO1695 | GCAATTTGCGGTGTTGCATG |
| **Genotyping primers** | | |
| *SALK T-DNA Left Border* | LB1.3 | ATTTTGCCGATTTCGGAAC |
| *GABI-KAT Left Border* | Gabi-KAT 03144 | ATATTGACCATCATACTCATTGC |
| *fc2-1 (GabiKat_766H08)* | For. JP283 | GAGCAACGCCAAACATAGAAG |
| | Rev. JP284 | TCAAAGGCAATGAATGTTTCC |
| *aos1-1/dde2* (*SALK_017756*) | For. WLO1805 | CGAGAAATTAACGGAGCTTCC |
| | Rev. WLO1806 | CTAACCGGAGGCTACCGTATC |
| *eds16-1/sid2-2* | For. WLO1803 | CCGAAAGACGACCTCGAGTT |
| | Rev. WLO1804 | TGGAGTTGGATGCAGAGCAG |
| *pub4-6* dCAPS genotyping: *Hpa*II digestion, wt = 97 bp, mutant = 121 bp | For. JP742 | TATTAGAGTAGTGTGAGTCAGG |
| | Rev. JP743 | GATCCAGTGATTGTGTCATCC |
